# Towards an information theory of quantitative genetics

**DOI:** 10.1101/811950

**Authors:** David J. Galas, James Kunert-Graf, Lisa Uechi, Nikita A. Sakhanenko

## Abstract

Quantitative genetics has evolved dramatically in the past century, and the proliferation of genetic data, in quantity as well as type, enables the characterization of complex interactions and mechanisms beyond the scope of its theoretical foundations. In this paper, we argue that revisiting the framework for analysis is important and we begin to lay the foundations of an alternative formulation of quantitative genetics based on information theory. Information theory can provide sensitive and unbiased measures of statistical dependencies among variables, and it provides a natural mathematical language for an alternative view of quantitative genetics. In previous work we examined the information content of discrete functions and applied this approach and methods to the analysis of genetic data. In this paper we present a framework built around a set of relationships that both unifies the information measures for the discrete functions and uses them to express key quantitative genetic relationships. Information theory measures of variable interdependency are used to identify significant interactions, and a general approach is described for inferring functional relationships within genotype and phenotype data. We present information-based measures of the genetic quantities: penetrance, heritability and degrees of statistical epistasis. Our scope here includes the consideration of both two- and three-variable dependencies and independently segregating variants, which captures additive effects, genetic interactions, and two phenotype pleiotropy. This formalism and the theoretical approach naturally applies to higher multi-variable interactions and complex dependencies, and can be adapted to account for population structure, linkage and non-randomly segregating markers. This paper thus focuses on presenting the initial groundwork for a full formulation of quantitative genetics based on information theory.

## 1. Introduction

The critical questions for understanding a genetic system, its functions, structure and complexity, lie in the actual dependencies among the system’s variables, both the phenotypes and genotypes as well as external factors. The phenotypes, of course, can range from highly specific cellular or molecular measures to broader, functional and organismal level phenotypes. The information architecture of the genetic system’s variables is at the heart of the dependency problem, and the difficulty of determining this architecture from data is significant for truly complex systems, which well describes many important genetic problems. These problems are inherent in the challenges of the past concerning the genetic explanation of complex traits, the notion of missing heritability and the complex effects of gene interaction. Quantitative genetics has evolved substantially over the 100 years since Fisher and Wright laid its foundations in these papers [1,2], for example. It has been pointed out repeatedly, however, that while their methods were powerful and innovative, there are some problems with the general approach and the tacit assumptions inherent in them [3,4]. It is not that the classical methods are not correct and powerful, but rather that there are unanticipated subtleties and tacit assumptions that are often not recognized. The proliferation of new data types calls for additional approaches and different mathematical descriptions, and since the logic of using the classical variance methods to infer genetic architecture is flawed [4], new approaches are needed for this reason as well. Nelson, Petterson and Carlborg have argued effectively that the Fisherian paradigm has reached its limits in the ability to deal with complex traits and modern genetic data [3]. Their summary, “… many of the current tools are adaptations of methods designed during the early days of quantitative genetics. The present analysis paradigm in quantitative genetics is at its limits in regard to unraveling complex traits and it is necessary to re-evaluate the direction that genetic research is taking for the field to realize its full potential,” is a clear call for new quantitative approaches. It is also true that, in spite of the innovative statistical approach in 1918, the Fisherian methods have often been misunderstood and/or misused in present quantitative genetics. Huang and Mackay have pointed out and clearly made the case that the genetic architecture of quantitative traits simply cannot be inferred from variance component analysis which has been applied for that purpose in many studies over the years [4]. The logic of this use of variance analysis is simply wrong because the underlying assumptions that would allow such inference do not generally hold. It is clear that genetic interactions, called epistasis in one common use of this term, have been implicated as essential for understanding complex traits [5-9]. Though it has also been challenged as being unimportant in evolution [10], this seems unlikely to us. Recent results have, on the contrary, strongly supported the importance of interactions in understanding complex traits [7, 35-38], including those in humans, and it would seem that evolution cannot escape such an influence. In addition, quantitative inference of interacting loci will likely be important for understanding polygenic risk scores which are currently being generated using non-interacting, largely additive, models.

Here we propose that information theory can provide the foundations of a new approach to quantitative genetics which focuses on the information content of the genome and the advantages of information theory, and we begin the process of building that foundation with this paper. It is not our position that present methods are faulty, but rather that it is likely that establishing a new approach and formulation will reveal new insights and provide methodologies because of the fundamentally different viewpoint. For example, the ability to detect two locus dependencies without significant single locus dependence extends the analysis power beyond the Genome-Wide Association Studies (GWAS) method. This extension is a natural feature of the information theory formulation.

The application of information theory to genetic problems actually has a long history. It begins with the surprising fact that Claude Shannon, the architect of information theory [11], actually wrote his PhD thesis in 1940 on “a new algebra of genetics [12,13], which addressed some key issues in population genetics at the time. In later work issues relating to the relationship between evolution and the statistics of population genetics were tackled using concepts from Shannon’s information theory [14-16].

Information theory, while originally directed at understanding communications quantitatively, has been very effective well outside of this original domain and has been applied widely to physical, biological, and chemical problems, and to other fields [21,17,18]. In almost all scientific domains the problem of inferring the quantitative dependencies among measurable variables, and even causal relationships, is the central problem, and the information measures, functionals of probability distributions, have been shown to be powerful tools in these problems of inference. We have previously shown how information theory methods can be used to analyze complex data, and also shown how genetic data is amenable to some such applications [19-21]. Here we extend both the formulation of the relationships and methods and their interpretation and recast the theory into a more comprehensive description of quantitative genetics. While we take only the first few steps here towards a full information theory of complex genetics, we show how this approach forms a fruitful way to describe the complex genetic architecture of a system. Specifically, we describe familiar concepts like gene interaction, pleiotropy, penetrance, degree of epistasis and heritability in terms of information theory.

Our concept of the information architecture of a system derives primarily from the idea of using information measures to define the levels of dependencies among variables. Information theory, being model-free, is broadly applied to extracting statistical properties from the data, which are in turn determined by the joint probability distributions of the variables.

Information measures have the advantage of being completely agnostic of any models or prior assumptions affecting dependencies, unlike many commonly used methods in genetics, particularly including correlation methods. This model-free character allows the data to fully drive the conclusions. These methods also reduce the sensitivity of the measures to small variations in the data, and to the limitation of small sample sizes. Thus, we argue here that the application of information theory to genetics can provide a powerful approach to deciphering the structure of complex genetic systems and to extracting their information architecture, which is distinct from the genetic, or model architecture. This paper advances our previous work in which we defined an information landscape [19] and illustrated the use of discrete functions and noise on this landscape to analyze genetic data. Here we focus on specifically elucidating the relationships of three-variable dependencies and complete this picture by providing a way to extract the specific functional nature of dependencies for variables whose dependency has been detected and measured.

General as it is, the application of information theory to any specific area carries with it certain assumptions and premises which need be made explicit. The principal caveats that must be addressed are these. The idea that the statistical inferences from the data reflect the subtle features of variable dependency assumes that the sampling issues and density of data represent these features in sufficient detail for information methods to make reliable estimates of the fundamental quantities, the entropies. In actual use this is often a rough approximation only and the approximation must be explicitly quantitated and its meaning acknowledged. We discuss this question later in the paper and for the purpose of explication initially simply assume for the moment that the data set is large enough to be fully reflective of the underlying relationships. It is also clear that by its nature information theory is inadequate to fully represent some distinctions among certain distributions. There are indeed distinctly different distributions with identical information measures. The mapping of probability distributions of variables into information measures is decidedly many-to-one. There are therefore several models and architectures that may have the same sets of measures. Another caveat depends on the question of how many variables participate in synergistic dependencies in a complex system since the number must be carefully controlled in any practical application because of statistical and computational limits [22]. While the method is entirely general, we limit ourselves in this paper to considering two and three variable dependencies only. This is sufficient to demonstrate the formulation and to illustrate its usefulness, and the power of the three variable method is amply demonstrated.

To make the formulation more self-contained we add a short primer on the key information theory quantities. First and foremost is the definition of Shannon’s entropy. For m possible states of a variable, *X*, {*x*_*i*_}, where the probability of a sample or subject *i* having a value of *x*_*i*_ is *p*_*i*_, the entropy of the variable *X* in this data set is 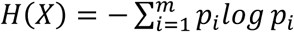. The joint entropy of two variables *X* and *Y* is defined in the same way where the possible states are those of the pairs {(*x*_*i*_, *y*_*j*_)}. The conditional entropies are obtained from the Shannon formula by simply using the appropriate conditional probabilities. An important measure that assesses the information in one variable about another is the mutual information. For two variables X and Y this is denoted I(X,Y), and is defined as *I*(*X, Y*) =H(X)+H(Y)-H(X,Y). If the two variables are independent of one another this is zero, as expected, since the joint entropy in this case is simply the sum of the two single entropies. The joint entropy can be extended to three variables by simply using the distribution of values of triplets of the variables, which are also obtained from the data in practice. All of the information measures used here can be expressed as sums and differences of entropies. In Appendix E we briefly address the important issue of estimation of entropies from data, the accuracy of which depends most sensitively on the amount of data available and the range of variable values considered. The errors in our calculation of entropies can be estimated, and must be kept in mind, but we rely here on the small variable alphabet sizes, and the large number of data samples to keep these small.

The symmetries of the relationships among the information functionals are surprisingly simple, but also subtle. The multiple measures of information theory have strikingly symmetric relations and number of symmetries that we have previously reported [23-26]. The symmetries all derive from the fact that all information measures are specific linear combinations of joint entropies, like the mutual information, organized by lattices whose partial order is determined by inclusion of variable subsets. In addition, there are a number of problems that can be fully analyzed for discrete functions, which are the most common manifestations of the variables we deal with in data analysis. By this we mean that the dependent variables in a complex system can be viewed as functions of one another, and the discrete values of the data can therefore be viewed as reflecting these discrete functions. While real genetic data has various levels of probabilistic determinants and “noise”, much of the character of the dependency can be represented by multi-valued discrete functions, which are mixed with various levels of “noise” to describe the realistic intervariable dependencies. This gives us a distinct mathematical advantage since, in principle, we can characterize the properties of all possible discrete functions with finite alphabets. We examine here the properties of discrete functions and their information architecture and relationships, show in detail how functions can be classified, and examine the extension of this analysis to include probability density functions that result from adding “noise” or subtracting determinism from the discrete functions.

## 2. Overview of Formalism

The complexity of genetics arises not only from the interactive functions encoded in the genome, and the range and complexity of phenotypes, but also from the structures of study populations and inheritance patterns in complex pedigrees. In this paper, while recognizing the important effects of population structures on quantitative genetics measures, we defer addressing these important issues so that we can restrict our considerations here to large, randomly mating populations, described as *panmictic*, recognizing that no natural population is fully panmictic, and few artificial, experimental populations are panmictic in practice. We will consider population structure issues in a later paper.

The basic components of the formalism presented here are summarized in these five points:

1. The information measure we call the *symmetric delta*, [21] as shown in section 4, is used to detect the dependence of subsets of loci with phenotypes in the data. In this paper, we consider pairwise and three-way dependencies.
2. The general relation between genetic loci and phenotypes is embodied in discrete valued *loci-phenotype tensors*: *f*(*X*_1_, *X*_2_, … *X*_*n*_), where {*X*_*i*_} is the set of *n* genetic loci and the function determines the phenotype. This is identical in two dimensions to what geneticists often embody in a matrix connecting three variables, called a “gene-phenotype table”. We limit ourselves to one or two genetic variables (loci) here. Without loss of generality we could include multiple phenotype variables as well.
3. The essential “noise” distributions, when added to these tensors, form the genotype-phenotype tensors (GPT) which describe the phenotype in terms of loci, noise and penetrance

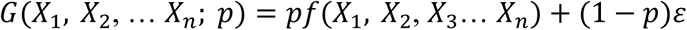

where *ε* is a “noise” function, and (1-*p*) is the noise level (*p* is the penetrance.) The noise can be assumed to follow a particular structure (*e.g.*, uniform random noise).
4. The tensor functions *f* and *G* for tuples of variables with significant dependence are inferred from the data using relatively simple algorithms.
5. These tensors are then used to calculate penetrance, heritability, gene interactions and pleiotropy.

This paper is organized as follows. We first present the basic discrete function expression of genetics, gene-phenotype tables which assumes full genetic dependence (with no “noise”), then review the basics of the information measures previously introduced [19-22]. We then describe some specifics of three variable dependencies and the symmetries that their information measures exhibit [26]. We review the information landscape notion we previously proposed [19] and extend it to a more general form. Introducing a formal tensor structure for extending the information landscape allows us to systematically handle all probability distributions, which are essential for the introduction of “noise,” for arbitrary size alphabets (possible discrete values of variables.) This formulation shows that the information content of the discrete functions is strongly dependent on both the alphabet size and the symmetries of the functions. This rich area is partially explored here but provides us some initial insights and a flexible set of theoretical tools with which to characterize complex genetic systems. We then define a set of transformations that maps the three-variable functions into a two-variable function space and allows us to greatly simplify the identification of the functional structure of the inferred dependencies.

We discuss the implications of these results and tools for the analysis of genetic data using information-based methods, and describe, in addition to penetrance, the genetic notions of gene interaction, pleiotropy, and heritability in terms of information theory measures. Finally, we apply our methods to some real yeast data and discuss the analysis of complex genetic data [33].

## 3. Discrete Functions and Genetics

The classic genotype-phenotype table for two loci can be usefully considered as a discrete function where the phenotype variable, *Z*, is expressed as a function of the two genotype variables, *X* and *Y*. Diploid binary variants for *X* and *Y* are, of course, three-valued, haploid binary variants are two-valued. While the phenotype alphabet can be any size, in principle, we also use three values for the phenotype alphabet (0,1,2). The alphabet can certainly be expanded to include more than binary allele variants, but for simplicity we do not consider these in this paper. Often a two-valued variable is sufficient to effectively describe a phenotype, but quantitative phenotypes require larger alphabets. These tables are similar to the Punnett squares in classical genetics. Consider the discrete functions where all three variables, *X, Y*, and *Z* are 3-valued, and *Z = f(X,Y)*, with *X* and *Y* independent. Each of the functional relationships can be represented by a 3-by-3 table. Table 1, for example, shows three functions that can be seen as tables for logical AND, logical XOR, and equality (EQ) functions, extended to three variables. In these tables the genotypes are encoded as: 0 = homozygous major alleles, 1 = heterozygote, and 2 = homozygous minor allele.

**Table 1.** Examples of extended (3-by-3) tables defining 3-valued versions of genetic functions. From left to right these correspond to logical AND, XOR, and EQ functions. These functions can be represented in a linear notation (by reading the tables left-to-right and top-to-bottom) as 000011012, 012120201, and 000111222, respectively.

|  |  |  |  |  |  |  |  |  |  |  |  |  |  |  |
| --- | --- | --- | --- | --- | --- | --- | --- | --- | --- | --- | --- | --- | --- | --- |
|  |  | X |  |  |  |  | X |  |  |  |  | X |  |  |
|  | AND | 0 | 1 | 2 |  | XOR | 0 | 1 | 2 |  | EQ | 0 | 1 | 2 |
| Y | 0 | 0 | 0 | 0 | Y | 0 | 0 | 1 | 2 | Y | 0 | 0 | 1 | 2 |
|  | 1 | 0 | 1 | 1 |  | 1 | 1 | 2 | 0 |  | 1 | 0 | 1 | 2 |
|  | 2 | 0 | 1 | 2 |  | 2 | 2 | 0 | 1 |  | 2 | 0 | 1 | 2 |

These discrete functions describe the phenotype as a function of the two genetic loci. The EQ function is a degenerate case for which *Z* is only a function of *X*. The general scheme can also be thought of as implementing a three-valued logic. We can call the function defined by the leftmost table (Table 1) an extended AND, because the lower value of the two arguments, X and Y, dominates, as in binary logic. Since there may be phenotype determinants other than these two loci, as described above, we will generally need a more complex function to describe the phenotype in a population. A three-variable tensor, which we consider in detail in a later section, can embody this complexity. The discrete function is generally modified by the random effects of both unknown and stochastic effects, the “noise” represented by a random function, and the penetrance, the degree of determination of the phenotype by the genetic variables.

The diploid case considered here is, of course, most commonly encountered in genetics of mammals, but the haploid case is not unknown in genetic data, for example in the case of recombinant inbred populations and organisms that can grow and divide as haploids. We apply our methods to an example of a haploid case in data from *Saccharomyces cerevisiae*. In the case of haploid genetics, the alphabet of values for the genetic variables is binary so that the genotype-phenotype table is 2×2 and the logic is essentially Boolean. This simplification can be very useful for practical calculations, as we discuss later.

## 4. Elements of the Theory

### 4.1 Genetic dependence relations

We begin by reviewing some definitions and previous results, and then introduce extensions of these relations. The first important point is that mutual information, an inherently pair-wise measure, is unable by itself to capture the full information in dependencies. Full representation requires many variable subsets, but even for three-variable tuples considered here mutual information is insufficient and requires additional measures to fully characterize the dependence among three variables. As has been pointed out before, a clear example is the exclusive OR relationship (XOR) for any size of alphabet [19]. For the binary alphabet, three-variable case it is evident that the mutual informations between all pairs of variables for this function vanish. We have demonstrated that the ternary XOR-like functions (Table 1) also exhibit this property [19]. It is also true for any size alphabet and is reflective of the symmetry of the dependencies.

Even the interdependency of two variables has a surprising level of complexity in the ways it can be expressed. Mutual information has several equivalent mathematical expressions. The most common form is as a difference of entropies, as described in the introduction. In terms of the conditional entropies we also have these symmetric expressions for mutual information.

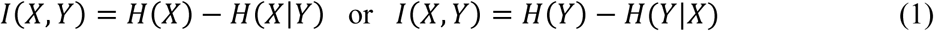

An important information measure is a generalization of mutual information for multiple variables, called the interaction information, or co-information [24,25]. For *n* variables this is defined by the recursion relation

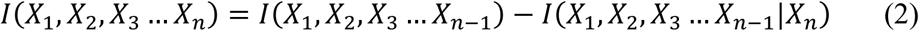

This measure can also be expressed by the sums and differences of joint entropies of the full set of variables *ν* = {*X*_1_, … *X*_*n*_}, (represented by Möbius function for the lattice of subsets, *τ*, in this formula)

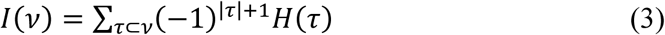

For three-variables the interaction information is simply expressed in terms of the entropies.

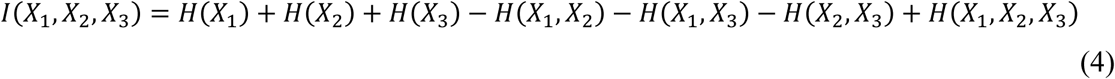

We define the *differential* interaction information (delta) as the change in the interaction information that occurs when we add another variable to the set. In general, where *X*_*m*_ ∉ *ν*_*m*_ and *ν*_*m*_ ∪ *X*_*m*_ = *ν*, the differential interaction information is defined as

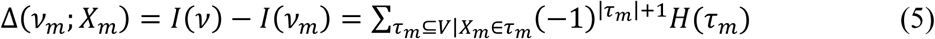

The general measure of the fully collective dependence among all variables, the symmetric Delta, 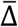, is defined as the product of deltas:

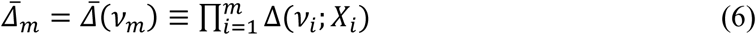

This is the measure we have previously proposed and used for measuring collective dependence of a set of variables. For three variables the differential interaction informations (the “deltas”) can be obtained by permutation of the variables.

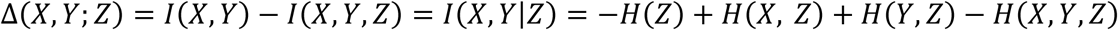

and the symmetric delta for three variables is

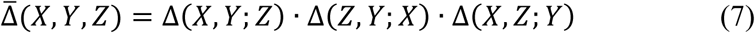

*H*(*X*_1_, … *X*_*n*_) into a sum of terms each of which depends on the number of variables, using the Möbius inversion [26]. This gives us an expression for the entropy as a sum over interaction-informations over all possible subsets of variables. This approach generates a series of approximations in the number of variables considered, and represents a practical, general and systematic way forward in the genetic formalism for more than three total variables, in that it provides the appropriate approximation for each limiting assumption. We will illustrate and use this approach in future work.

#### 4.1.1 Multi-information as total dependence

Another important information measure that we will use in several ways is the multi-information for *n* variables (originally defined and called “total correlation”, by Watanabe [27] and discussed and used by many others [28,29]). It is defined as the difference between the sum of entropies of each variable separately and the joint entropy of all the variables together:

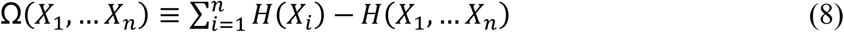

The multi-information is essentially the collective measure of all dependencies among the *n* variables; that is, the sum of dependencies for all possible subsets of variables. It is zero only when <u>all the variables</u> are independent, so it does not distinguish among the orders of dependency. This stands in contrast to the symmetric delta, which is the measure of the full, synergistic dependency of all the *n* variables together. It is zero when <u>any one of the variables</u> is independent of the others. Since the multi-information deals with dependence of all possible subsets, and the symmetric delta deals with dependence of the entire set, they are like bookends of the dependency measures. As shown in the next section the multi-information is a key element in the quantitative relationships we use in this formalism.

#### 4.1.2 Three-variable dependencies

While the restriction to pairwise dependency analysis is equivalent in concept to classical association studies in genetics, sufficient for some problems, the detection of even three-variable dependencies can add much to the power of the analysis and is essential for any genetic system that involves pleiotropy or gene interaction. Note that pleiotropy is defined as the dependence of one genetic variant variable and two phenotypic variables. We focus in this section on understanding the key relations for systems at the three-variable level. The relations among the three-variable information measures are simple, but subtle, and illustrate the strong symmetries inherent to the information measures. Furthermore, it is useful to examine carefully the bounds on their values. But first, a few more preliminaries.

From here on we will use a simplified notation, where the three variables are labeled by integers: *X* → 1, *Y* → 2, *Z* → 3. Wherever the meaning is clear, we will abbreviate using these labels within a subscript; for example, H(X, Y, Z) → H_123_. The relations between the mutual informations and the multi-information, and the deltas (where we define the notation Δ_1_ ≡ *I*(2,3|1), Δ_2_ ≡ *I*(1,3|2), Δ_3_ ≡ *I*(1,2|3)) are provided by these equations:

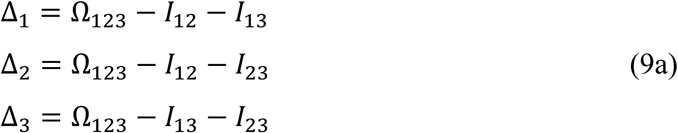

We derived these previously [19], but they are easily shown to be true by simply expressing the measures Δ, Ω, *and I* in terms of sums and differences of entropies. Since Ω always refers to all three variables we can drop the subscripts for this quantity without ambiguity in most cases. For two variables only, of course, Ω = *I*_*ij*_, the mutual information. It can easily be seen that these equations are symmetric in the variables, and the only asymmetry arises from the differences among these terms. The above relations for three variable dependencies can, of course, also be formulated conveniently in matrix form, which we show in Appendix C. This matrix equation may be a useful tool for further exploration of three-way dependence symmetries.

For genetic data, where X and Y are independently segregating genetic loci, valid for panmictic populations, and Z is the phenotype variable, the three mutual informations in (9a) become two since *I*(*X, Y*) = *I*_12_ = 0. The assumption of independently segregating variants is essentially equivalent to assuming linkage equilibrium. In this case there are only three relevant measures in the set of relations (9a), Ω, *I*_13_, and *I*_23_, and the relationship is significantly simplified.

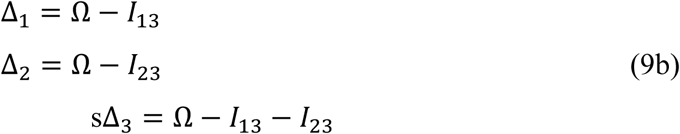

We can normalize the Equations 9b by dividing through by Ω, the total of all dependencies, as long as there is some dependency so that Ω > 0. We get the normalized delta coordinates (only for the case of *I*_12_ = 0) which were the coordinates used in [19] to define the geometry of the information landscape. The coordinates of the information landscape are these:

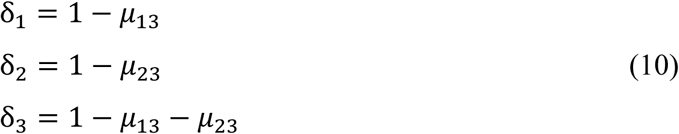

where the δ′s are the normalized Δ′s, and the *μ*^′^*s* are the normalized mutual informations. We can rearrange the above equations into a simple relation for δ_3_ as a function of δ_1_ and δ_2_:

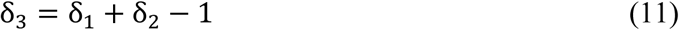

The condition for δ_3_ to be non-zero then is δ_1_ + δ_2_ > 1. This is one side of the line defined by δ_1_ + δ_2_ = 1. Let us look more closely at the constraints on δ_1_ and δ_2_ imposed by *I*_12_ = 0. If we look at the 3-D space defined by the three δ’s, which is what we call the information landscape, we can see that we have three coordinates and one linear constraint that thereby defines a two-dimensional plane. One natural question regarding bounds of the landscape is whether negative coordinates are possible. The answer is that they are not. The key inequalities that bound these quantities are intuitive and elementary but still not entirely obvious and we state them explicitly and present the proofs in Appendix A.

The interaction information, *I*_123_, is defined in terms of the entropies [24,25] as

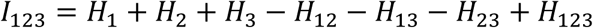

and by the definitions of mutual information we have

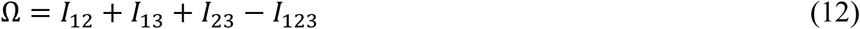

Notice that if *I*_12_ = 0, by proposition 2 this expression implies that *I*_123_ ≤ 0.

A few more points about dependencies among genetic variables are in order here. Equation 9b applies when *I*_12_ is strictly zero, however *I*_12_ may as well be nonzero in real data, because of disequilibrium or because of noise in the data, including sampling induced fluctuations. We will deal with the linkage disequilibrium (LD) issue in a future paper, but it is important to note that even in the presence of LD the symmetric delta represents the full interaction score for any triplet, including the contribution due to LD. A significant problem to be discussed in a future publication is that it is more difficult in this case to extract the quantitative score for the strictly three-way component (we call this the epistatic component). The potential entanglement of epistasis and linkage disequilibrium, which is often overlooked in genetic analyses, is at the heart of this issue.

There are many ways of expressing the set of relationships described above for three variables. For example, Equation 9 leads directly to the expression for the multi-information as in Equation 14. Since it is clear that if the dependencies are pairwise, and *I*_12_ = 0, then the mutual informations contain all the dependence, in which case Ω = *I*_13_ + *I*_23_. Thus, in this case for three-way dependence, we can ascribe the epistatic component (three-way) to the value of −*I*_123_ (the minus sign comes from our sign convention above). In other words, in the case of linkage equilibrium the interaction information is the epistatic dependence measure. This is a useful way to decompose the multi-information. This relation for the triplet dependencies is illustrated in Figure 1. We emphasize again that the epistatic component is −*I*_123_ only when *I*_12_ = 0. In the general case −*I*_123_ is equal to the epistatic component minus the information shared by 1 and 2 affecting 3. The above equations allow us to define several important limiting conditions. This is further illustrated in Figure 2. We can summarize these constraints on the basic measures and their implications or interpretations simply and this is presented in Table 2.

**Table 2.**
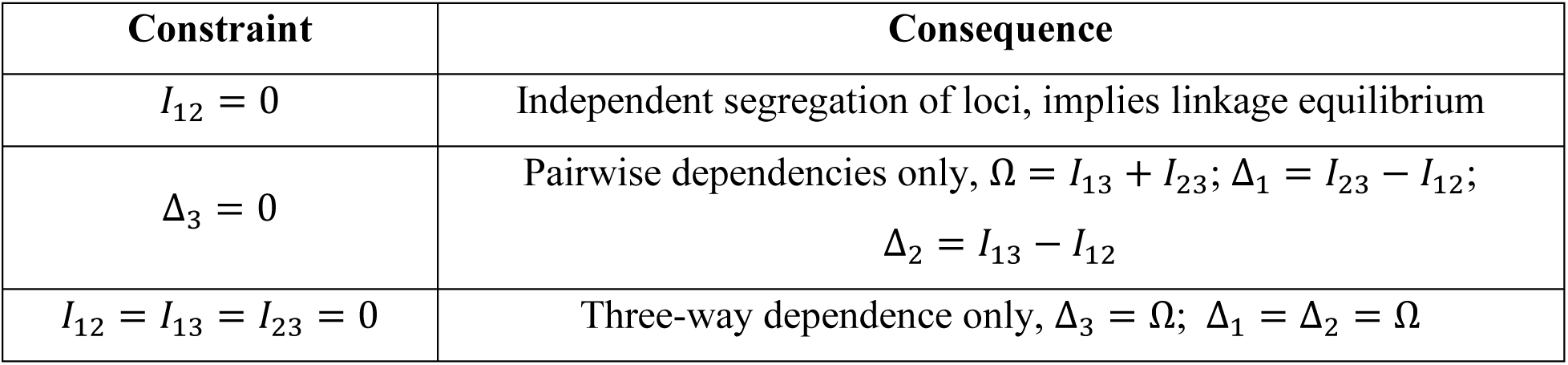
Several limiting constraints on the information relations, with their interpretations or consequences. Keep in mind that these rules apply strictly only to the discrete functions without noise.

| Constraint | Consequence |
| --- | --- |
| $I_{12} = 0$ | Independent segregation of loci, implies linkage equilibrium |
| $\Delta_3 = 0$ | Pairwise dependencies only, $\Omega = I_{13} + I_{23}$; $\Delta_1 = I_{23} - I_{12}$;<br>$\Delta_2 = I_{13} - I_{12}$ |
| $I_{12} = I_{13} = I_{23} = 0$ | Three-way dependence only, $\Delta_3 = \Omega$; $\Delta_1 = \Delta_2 = \Omega$ |

**Figure 1.**
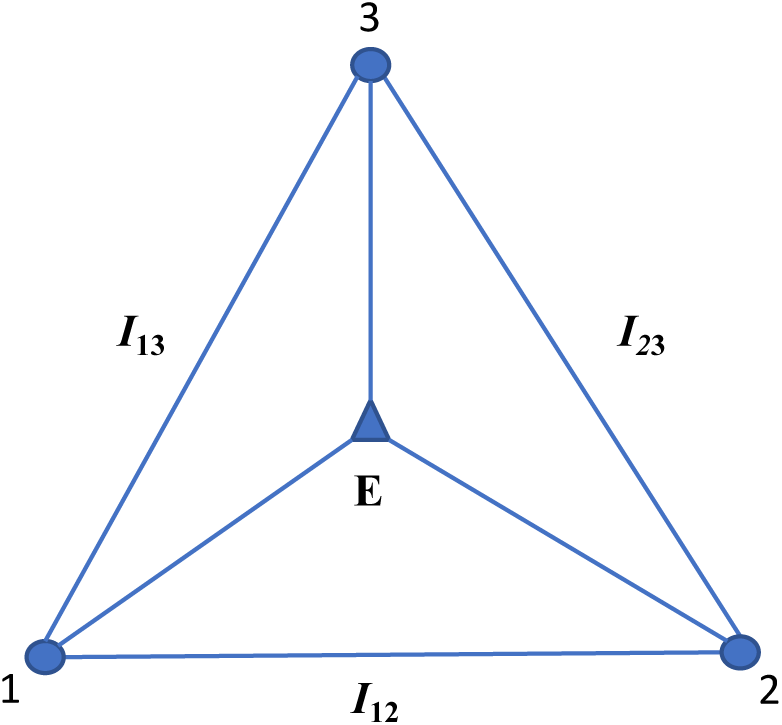
Three-variable dependencies that make up the multi-information or total correlation. (we adopt the convention here that X is 1, Y is 2 and Z is 3). The lines represent the components of dependence among the variables (small circles) as in the above equation, where the epistatic component is represented by the lines emanating from the triangle. The epistatic component is *E* = −*I*_123_ + *S*.

**Figure 2.**
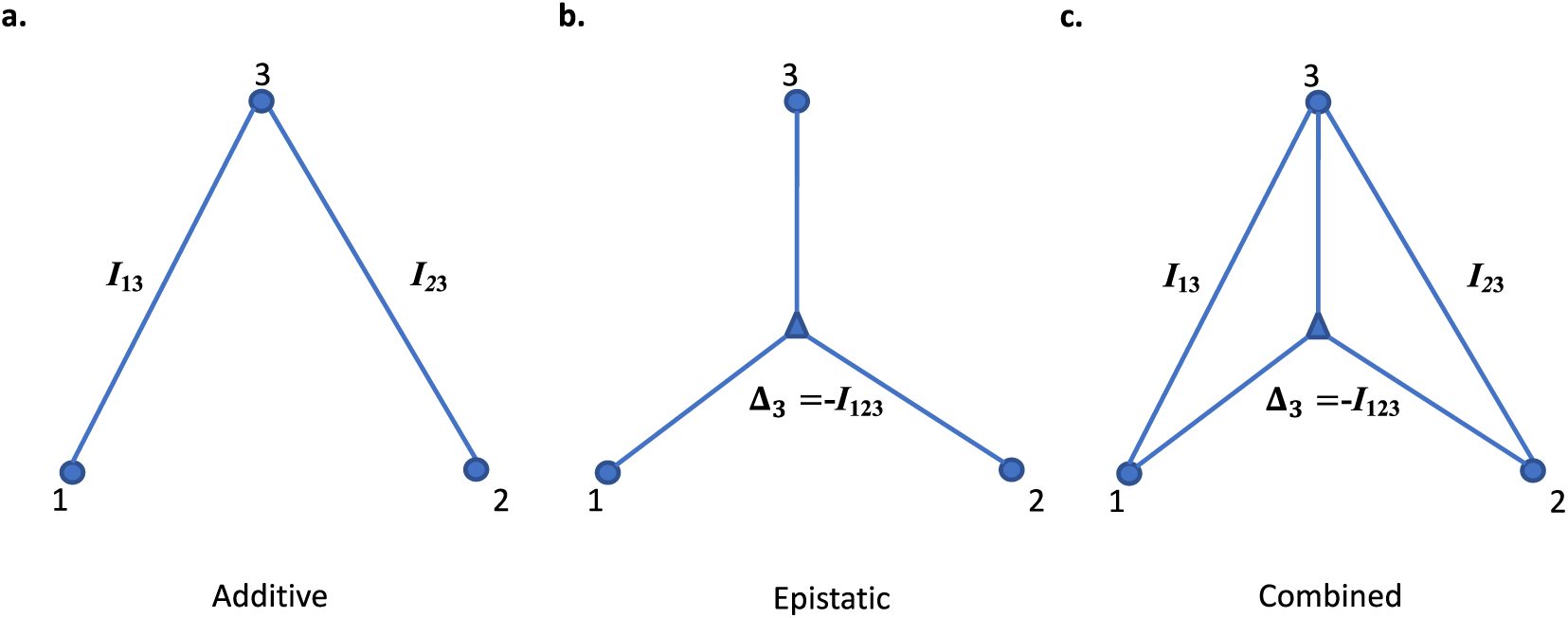
Independent segregation interaction relationships. The genetic contributions, (1 & 2) to the phenotype, 3, illustrating the distinction between the additive and epistatic effects.

#### 4.1.3 The components of genetic dependency and their measures

The genetic architecture of a phenotype is determined by the dependencies among the genetic variables and the phenotype variable. The application of the information formalism can, however, be rather subtle and care must be taken in its interpretation. In this section we define the problem in a bit more detail and make the specific connections between information theory quantities and genetic quantities. The dependencies of phenotypes on more than one genetic locus define what we mean by genetic interactions and consists of a wide, but finite, range of possible forms of interactions. These effects have been recognized for 110 years when William Bateson proposed this as an explanation for deviations from simple Mendelian ratios. If a phenotype is dependent on two loci, each exclusively in a pairwise fashion, we call this effect additive, and distinguish it from what in the usual terminology is called an epistatic effect or interaction. Fisher called this statistical epistasis *epistacy* and attributed the deviations from additivity to his linear statistical model. Many modern authors have argued and provided evidence that gene-gene interactions are rather common [9]. The most common way to deal with these interactions quantitatively, however, has been to use regression methods [30], and more recently, other machine learning tools. In all these cases, however, the starting loci are most often those identified by GWAS or some pairwise method, which will then miss those loci that are invisible to pairwise methods.

Quantitating “gene interaction”, that is, measuring the amount of the phenotype that depends on the combined markers, can be done naturally with the measures defined here. We need to be precise, however, in defining what we mean by gene interaction, and we need to distinguish additive effects from epistatic interactions, the former being strictly pairwise, the latter not including any pairwise effects. Again, we are here assuming independently segregating variants and *I*_12_ = 0.

If the genetic variant variables are *X* and *Y*, and the phenotype variable is *Z*, we consider all possible three variable dependencies, as in Figure 1. In this general, three-variable case we can quantitate the information contribution of *X* and *Y* to the determination of *Z* by the mutual information between Z and the joint X,Y variables, *I*(*Z*, (*X, Y*)). Using the mutual information chain rule

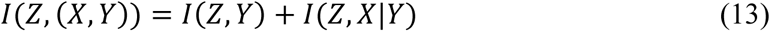

and identifying *I*(*Z, X*|*Y*) = Δ_*Y*_ and using Equation 9a we have simply

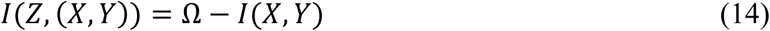

In the case of independent segregation of markers, where *I*(*X, Y*) = 0, this becomes *I*(*Z*, (*X, Y*)) = Ω, as expected since in the absence of shared information between X and Y the mutual information *I*(*Z*, (*X, Y*)) describes the full extant dependence. As shown in the previous section the decomposition of the information contributions becomes simple in this case. In Figure 2 we illustrate the nature of the dependencies.

We wish to emphasize that the relationships present here permit the decomposition of the information structure, the dependencies, of the variables. It is important that the dependency can be decomposed and we can determine what fraction of the dependence is pairwise and what fraction is three-way dependent (synergistic or epistatic). These fractions can be derived simply from the equation Δ_3_ = Ω_123_ − *I*_13_ − *I*_23_ provided that *I*_12_ = 0. Since the pairwise dependence of the phenotype on the two loci is the sum of the mutual informations, *I*_13_ and *I*_23_, and the total dependence is Ω_123_, the three-way dependence is given by their difference, Δ_3_. The fractional dependencies are then simply the ratios

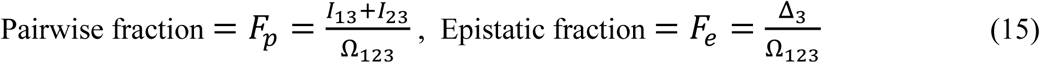

If the genetic variables are 1 and 2, and the phenotype is 3, these fractions represent the pairwise additive contributions of 1 & 2 to the phenotype, *F*_*p*_, and the non-additive, or epistatic, contribution, *F*_*e*_.

The Equations 14 and 15 apply in this case and the separation of the additive and non-additive, or epistatic effects is clear.^1^ We will address the more complex case of non-zero linkage disequilibrium and related effects in a future paper.

The epistatic interaction in the case of no disequilibrium is measured entirely by Δ_3_. This is also rather intuitive since the multi-information, Ω_123_, quantitates the total dependence and the mutual information quantitates the pairwise dependencies between each variant and the phenotype. Thus, their difference measures epistatic gene interaction.

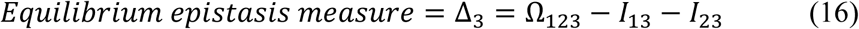

There is another kind of three-variable dependence that is important in genetics. A single genetic locus affecting two distinct phenotypes, which is called *pleiotropy*, can be described by the general equations, but the limiting constraint of independent segregation which makes the mutual information between variants vanish does not apply in this case. The analogous constraint, however, is that the mutual information between phenotypes vanishes. We consider pleiotropy briefly in our discussion of the yeast data. Full pleiotropic analysis can be rather complex, however. For example, unlike for two genetic variables, we cannot easily understand what it means to decompose the information contributions of one genetic variant and phenotype on a second phenotypic variable. The potential complexities are both interesting and significant and will be considered in future work.

### 4.2 Information theoretic relations and symmetries

When two loci (*X, Y*) are involved in determining a phenotype, *Z*, we can represent the relation as a genotype-phenotype matrix. These three-variable matrices have discrete values and thus are discrete functions of two variables, *Z*(*X,Y*). We have shown that the three-dimensional information landscape, defined by the three normalized deltas from Equation 10] is a plane when *I*_12_ = 0, and under this condition all discrete functions lie on this plane.

#### 4.2.1 Discrete functions

There are several possible ways of defining the information content of discrete functions, and discrete functions are a useful way to characterize quantitative genetic relations. The usual genotype-phenotype tables for two genetic loci used in classical genetics are just this kind of discrete function and therefore the information in these functions is the key to quantitative analysis. Here we define the inherent information as the measures calculated from the probabilities inferred directly from the function. Note that all discrete functions map into distributions, but not all distributions are discrete functions. Information measures (all are linear combinations of joint entropies) are functionals of distributions. However, the finite size of the set of discrete functions (for a given set of alphabet sizes) and the (infinite) size of the set of all possible distributions are incommensurate. There are a finite number of discrete functions for any finite number of variables and alphabets, but there is an infinity of distributions for any finite number of variables and alphabets. The addition of “noise” to the discrete functions generates an infinite range of distributions. As we will see this is a key consideration in quantitative genetics. The “noise” determines the penetrance of the genetic dependence on the discrete function.

As shown in [19], we can map all the discrete functions onto the information plane (for example, there are 19,683 functions on this plane for the 3×3 case). When the information measures are calculated for the 3×3 functions and plotted in the plane they form simple rectilinear patterns for each value of Ω. The positions of all function families (those functions with identical normalized delta coordinates) are shown in Figure 3a.

**Figure 3.**
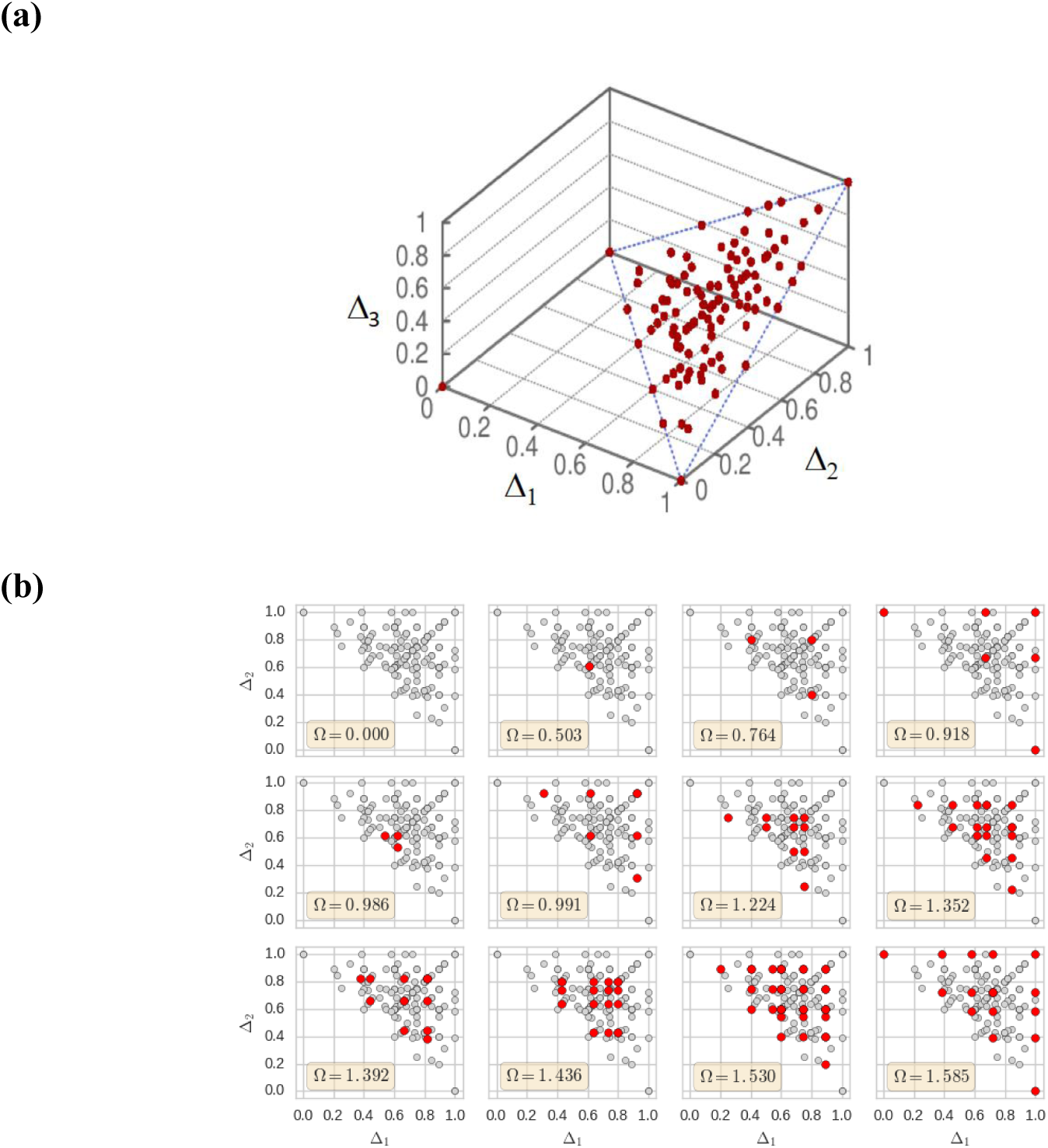
Function classes (3×3) on the landscape. Each spot in both panels represents a function class, or family. (**a**) The information landscape shows the orientation of the plane with respect to the 3D landscape. (**b**) A set of 12 panels, one each for the complete set of possible values of the multi-information, Ω, for the 3×3 functions. The plane is the projected diagonal plane of the three-dimensional landscape, the grey spots are the same for each panel and show the positions of <u>all</u> of the families of functions. The red spots are the families specific for each specific value of Ω. The upper left panel has no function as the information content of the uniform functions are zero, and all Δ′s are zero.

In Figure 3b the families are shown in different panels for different values of Ω, total dependence. Even though all functions in a family have the same delta coordinates, not all of the functions in a family need have the same value of Ω. Notice the symmetry in the triangular plane that results from the exchange of *X* and *Y*.

In the case of haploid genetics, the information plane for three variables shows a similar geometric symmetry, but with many fewer functions. Many published yeast genetic data sets are haploid, including the data we have analyzed here to demonstrate our methods [33]. Haploid genetic state variants are binary and since there are *N*^4^ discrete functions, where *N* is the alphabet size for the phenotype, *Z*, this can lead to a significant simplification of the information landscape for binary phenotypes. For *N* = 2 there are only 16 functions in all, but as *N* increases from 3 to 5 the number of functions grows rapidly, and there are 8 families of functions, each family having identical information coordinates. As the phenotype alphabet size, *N*, increases past 5 the number of families stays the same even as the number of functions grows rapidly. In Figure 4 the information plane and the families are shown.

**Figure 4.**
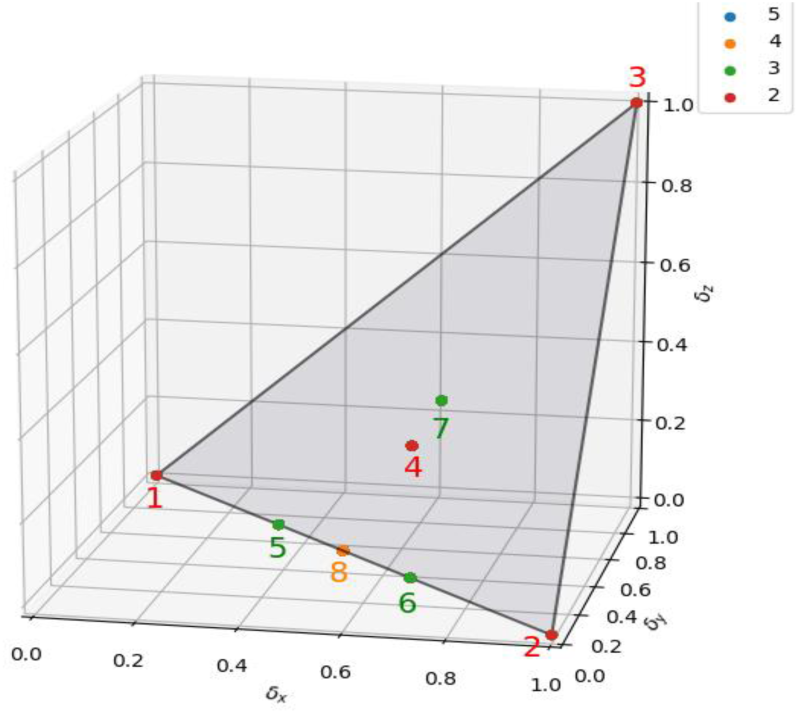
The information plane for haploid genetics, binary genetic variables. The color-coded points show the locations of the function families corresponding to the alphabet size of the phenotype, as indicated in the legend panel at the top right. Families 1-4 correspond to a binary phenotype alphabet. Families 5-7 are added for a three-letter alphabet, and family 8 is added for a four-letter alphabet. The blue dot in the legend is not seen since does not correspond to any specific family. The five-letter alphabet functions all fall into the previous eight families. While the limit is eight families, as the alphabet size increases the number of functions in every family grows. The families 1, 2, 5, 6 and 8 are functions with only pairwise interactions (*δ*_*Z*_0).

Finally, since we propose to use the symmetric delta of Equation 7a to find three-way dependencies it is natural to ask if the total dependence were equal for multiple triplets which discrete function would maximize the symmetric delta? As we show in Appendix B the answer is that it is the triplet with no pairwise dependence and only three-way dependence, the XOR-like functions. This is interesting in several respects, first because having the product of the conditional mutual informations maximal for discrete functions where pairwise dependence vanishes seems unexpected, but more importantly because a distribution of triplet symmetric delta scores, maximizing at the XOR-like functions is a very useful indicator of specific functional dependence.

## 5. Inferring the Functional Dependence of Phenotype on Genotype

### 5.1 The meaning of “noise” in genetic data

There will always be “noise” in the data, which arises from two classes of sources, unknown variables and stochastic processes inherent to the biology and data acquisition processes. The word is in quotes here to emphasize the composite and subtle nature of the several factors that determine what “noise” is. This quantity can therefore only be inferred from the data when we can explicitly define the character and degree of the dependencies we are including. If we only consider direct, pairwise effects, for example, from each of two loci on a phenotype, then the interaction between these loci affecting a phenotype (what we call a three-way dependency) as well as any other more complex interactions will contribute to the “noise”. Likewise, if we only include the effects of the genetic variants that we have ascertained, the loci not included will contribute to the “noise.” Any variables or effects not included can potentially contribute to the “noise”. All unknown genetic variants and all other unknown environmental factors may contribute to what we call the “noise” in this point of view. In this way we both more tightly define the quantitative nature of genetic penetrance and also provide a well-defined method for a data driven estimate of the key quantities. We therefore have two fundamental steps in a general method for the inference of the relevant dependencies: first, the detection of levels of dependencies using the information theory measures, followed by the inference of the functional nature of these dependencies and the “noise” level. The “noise” plays a direct role in determining the penetrance of genetic effects.

### 5.2 Probabilistic tensor model

The discrete functions of three variables, interpreted as distributions are illustrated in the above landscapes (see Figures 3 and 4), where the information measures are calculated from these functions. Since any phenotype is not fully determined by genetic functions, “noise” is recognized as an important factor in quantitative genetics, as we emphasized above. What we mean specifically by noise, however, includes unknown sources of effect, as well as truly stochastic factors, both biological and technical. The mathematical noise function we use here, *ε*, thus arises from several sources, specifically including these six:

1. Measurement errors in any of the variables, both phenotypes and genotypes
2. Environmental influences on the phenotypes
3. Epigenetic effects
4. Stochastic developmental & physiological effects
5. The effects of uncharacterized genetic variants (rare SNPs, CNV’s, etc. - anything not included in our genetic variables), interactions that involve more than 2 genetic loci, which we do not include here, and weak effects from other two and three-variable tuples that are below a statistical threshold for consideration. These are all what are usually called genetic background effects.
6. Sampling noise (allele frequencies *vs* subjects, etc.), purely statistical fluctuations.

The “noise” as defined here, of course, is actually not noise in the usual sense of the word, but the composite of all unknown influences as well as truly stochastic inputs.

The discrete functions represent a vanishingly small fraction of all possible information functions. However, they can be used effectively to describe real genetic effects, and generalized, by adding a noise function that modifies the probability of occurrence of each possible alphabet value of the phenotype. This allows us to flexibly represent general distributions for any specific alphabet size, and thereby defines the “noise” in our functions as described above. It is clear that the locations of these general functions on the information landscape are continuously distributed, as illustrated in [19] where we introduced random noise into the discrete functions. Here we introduce a systematic formulation, combining discrete functions with noise, by defining tensor functions.

In general terms, the relation between genetic loci and phenotypes is embodied in this formalism in discrete valued *loci-phenotype tensors*: *f*(*X*_1_, *X*_2_, … *X*_*n*_; *Φ*), where {*X*_*i*_} is the set of *n* genetic loci and *Φ* is the phenotype. For two genetic loci this is identical to a “gene-phenotype table”. When we take account of “noise” distributions we add a uniform distribution to these tensors and form the genotype-phenotype tensors (GPT). The relation between these for *n* variables is simply

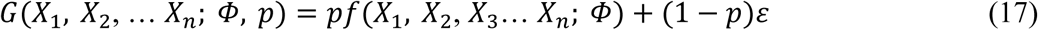

where *ε* is a uniform random “noise” function, and (1-*p*) is the “noise” level. The parameter *p* is the *penetrance*.

Since in the three-letter alphabet the discrete functions determine a third variable as a function of two others, the functions can be represented by 3×3×3 tensors, *G*_*ijk*_ where the indices *i* and *j* specify the genetic variables, and the index *k* specifies the alphabet of the phenotype. The entries in the tensor for a specific function define the probability for a position to have a given letter of the alphabet. Let us define *G*_*ijk*_ such that *G*_*ijk*_= 1 when *f*_*ij*_ = *k*, for the indices ranging over the alphabet, which are considered as non-negative integers here. Using this representation we can add noise to functions by entering probability values in the tensor that are other than 0 or 1. We use the uniform distribution, for which all letters are equally likely, as it is the only entirely unbiased representation of the “noise” component of the tensor.

For the genotype-phenotype tensor the fractional balance between the “noise” and the discrete function is a variable factor we call penetrance, in keeping with the usual use of the term in genetics. If the penetrance is 1 there is no confounding noise, and if it is small the genetic function plays that correspondingly small role in determining the phenotype. Note that if the penetrance is small the significance of the genetic effect is also small. Thus, there is a clear relation between penetrance and the p-value of the effect. This relation will be explicated further elsewhere.

We assume here that a full penetrance effect can be described by a single discrete function. It is possible that some linear combinations of discrete functions could be useful in some cases. We do not consider this more complex extension further in this paper.

It is important to see what happens to the coordinates as the penetrance decreases (“noise” increases). To see what the delta coordinates in the information landscape for tensor functions with low penetrance we examine the limiting ratios of the information functions. Since we cannot calculate deltas for the uniform distribution consider distributions infinitesimally close to the uniform distribution or close to zero penetrance.

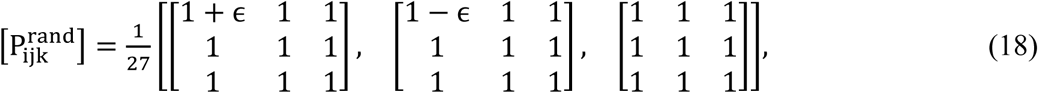

and calculate the delta coordinates in the limit *ϵ* → 0. The above tensor can be used to calculate the corresponding joint entropy, for example, for the 3×3 case.

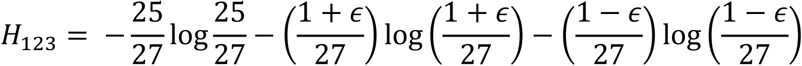

The delta-coordinates can be calculated from these entropies. The first delta-coordinate is

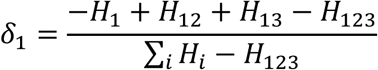

However, both the numerator and denominator of this expression go to zero in the limit when *ϵ* → 0, so we take the limit using L’Hospital’s rule. The first derivatives are also each zero, but using the second derivatives yields the limit:

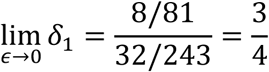

This is in agreement with our previous numerical results [19]. This location, 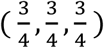, on the information plane has very particular properties that need to be carefully considered. When the tensor is completely dominated by the uniform probability the “noise” completely swamps out the information content of the functions, and the genetic information has no effect on the phenotype. This point corresponds to a value of the penetrance, *p*, of zero. It is the location on the information plane that we called the “black hole” previously [19]. As noise increases the functions all move their positions on the landscape, and they eventually converging on this spot.

### 5.3 An algorithm for inferring genotype-phenotype tensors

Since the relation between genetic loci and phenotypes is described by discrete valued *loci-phenotype tensors*, (see Equation 17), once we have used the information measures to determine that there is a significant dependence for a given set of variables, we need to infer the function itself to understand what the data implies. However, since the tensor is not described by a discrete function alone, we also need to infer the level of the essential “noise” distribution. As described above, together these components, the discrete function and the “noise” level, described by the penetrance, form the genotype-phenotype tensors (GPT), where the function *ε* is a uniform random “noise” function, and (1-*p*) is the noise level, and the parameter *p* is the penetrance (Equation 17). We will henceforth write the tensors using indices that range over the variables and the alphabets. Thus *f*(*X*_1_, *X*_2_, *X*_3_… *X*_*n*_, *k*) is written as 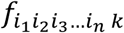, where the {*i*_1_} are genetic variant indices, and k is an alphabet index.

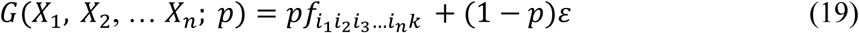

Given a data set and a significant tuple of variables (the dependence to be analyzed) there is a simple way to infer the function and *p.* The problem can be posed as follows for a three-way dependence. Let us represent the data by the data frequency tensor for a phenotype as a function of two genetic variants

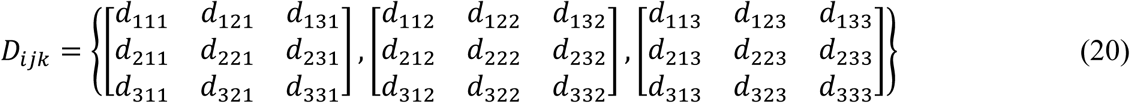

where all variables range over three-letter alphabets. This frequency tensor is defined to be normalized so that the sum of all components is 1.

Let us assume for the moment for simplicity of explication that the allele frequencies are equal. We will modify the resulting simple algorithm for non-equal allele frequencies later (note that this is moot for the haploid case we analyze in the last section). A simple, greedy algorithm for finding the most likely function, *f*_ijk_, from the data *d*_*ijk*_, simply identifies the maximum *d*_*ijk*_ for each letter of the alphabet *k*, and assigns a probability of one to that *k* and zeros to the other two for all *i* and *j*:

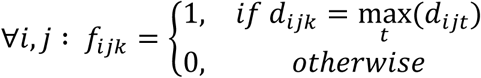

The algorithm is “greedy” in the sense that it takes the largest value of *d*_*ijk*_ for each *k* and gives it a value of 1. This prescription is incomplete, however, in that the maximum within each value of k matrix may not be unique. In this case we can choose the element to assign *f*_*ijk*_ = 1 randomly among the multiple maxima. We have not explored the quantitative impact of this source of noise but have found that for the large data sets explored so far, for example, the yeast data set in section 6, that there are unique maxima. This is another case where the more samples in a population the more frequently the noise is suppressed. The estimate of *p* in Equation 19 is the average frequency of the tensor elements not assigned a value of 1 in *f*_ijk_. If the expectation is taken over all tensor elements, since there are 9 non-zero entries for *f*_*ijk*_, we can write the expression for the penetrance, *p*, as

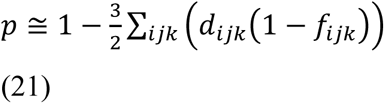

The algorithm yields the genotype-phenotype tensor

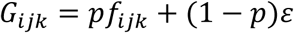

There are many ways of characterizing the resulting fit – measuring how well the data is described by such an inferred tensor function. In the spirit of the current formalism we can calculate the Kullback-Leibler divergence between *D*_*ijk*_ and *G*_*ijk*_, but a chi-squared test also works. Note that these tensors are normalized so that they can be treated directly as distributions.

### 5.4 The effect of allele frequencies

In the previous section we made the simplifying assumption that the allele frequencies were equal in order to more clearly explain the process. The frequencies are, of course, hardly ever equal. In order to deal with this issue, we can make a simple linear transformation of the matrices of the tensor to account for unequal allele frequencies, which slightly complicates the algorithm, but is not a fundamental difference. More importantly, it is essential to note that allele frequency differences can have strong effects on all of the information measures. Among other difficulties, strong interactions between loci, three-way effects, can potentially be masked by the rarity of key alleles in these loci. There is little that can be done to avoid this problem if it fully masks the interaction signal, however, the detection of weak interactions should therefore be looked at carefully in the population to ascertain whether the allele frequencies are involved in determining the strength of the signal. Additional cautions can include segregating the sample population to focus on more genetically homogenous subpopulations. This can only help, of course, if the numbers are large enough to properly analyze a subpopulation, but it may be a necessary step to avoid missing significant effects.

### 5.5 A simplification: Transformations of 3-functions into 2-functions

Each of the discrete functions of two variables, *Z*(*X,Y*), can be specifically transformed into a pairwise function without loss of dependency information. By this we mean expressing it as a function of a single variable that maps to values of the pair X and Y. For example, the 3×3 function we call an XNOR-like function is represented by this matrix that defines Z values (the columns are X, the rows are Y).

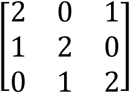

A transformation that maps pairs of (X,Y) values into another variable, call it W, can represent the essential information in the function: {(*X*_*i*_, *Y*_*j*_)} ⇒ *W*. What this means is that the matrix can be fully reconstructed by the mapping

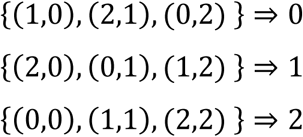

since this transformation simply yields values of the variable *W* such that *W=Z*. Every function can be mapped similarly. The resulting two-variable functions have the properties that the mutual information between W and Z are maximal. These transforms take these three-way functions with into a space of pairwise-functions with only pairwise dependencies. An important question, however, is whether the mapping of the three variable-function space into a pairwise-function space of the same size exhibits symmetries and redundancies that reduce the complexity of the one-to-one transformations. In other words, are there a set of “basis” transforms that can distinguish each of the 3-functions from one another when mapped into the pairwise-function space? The XOR and XNOR functions for three-variables have no pairwise dependence at all, and point out their importance for data analysis. In Appendix B we show in the three variable case that the symmetric delta is maximized by this family of functions.

The information measures described here are sensitive to dependencies, but do not define the functional nature of the dependencies, the functions themselves. In other words, the mapping of functions into delta coordinates is many to one, as is evident from the multiple functions in a family of functions with the same coordinates. The geometry of the information landscape, however, can usefully limit the possibilities since the delta values define a location in the landscape and thereby restrict the functions that could be generating the observed dependence. The detection of dependence, localization on the information landscape, followed by the identification of the actual function that then leads to transformations from 3-function space to 2-function space are the steps in the process of complete characterization of genetic phenomena with two-loci functions affecting phenotypes. The next question then, is how can this paradigm best be implemented?

### 5.6 Inferring the genotype-phenotype function and penetrance in simulated data

To test the effectiveness of the simple algorithm described in the previous section we created a simulated data set. We generated simulated data for 100 subjects. As test case, we used a specific discrete function and added a uniform “noise” function. The discrete function of three variables chosen in the case described here is shown in Figure 5. This function exhibits both pairwise and three-way dependence.

**Figure 5.**
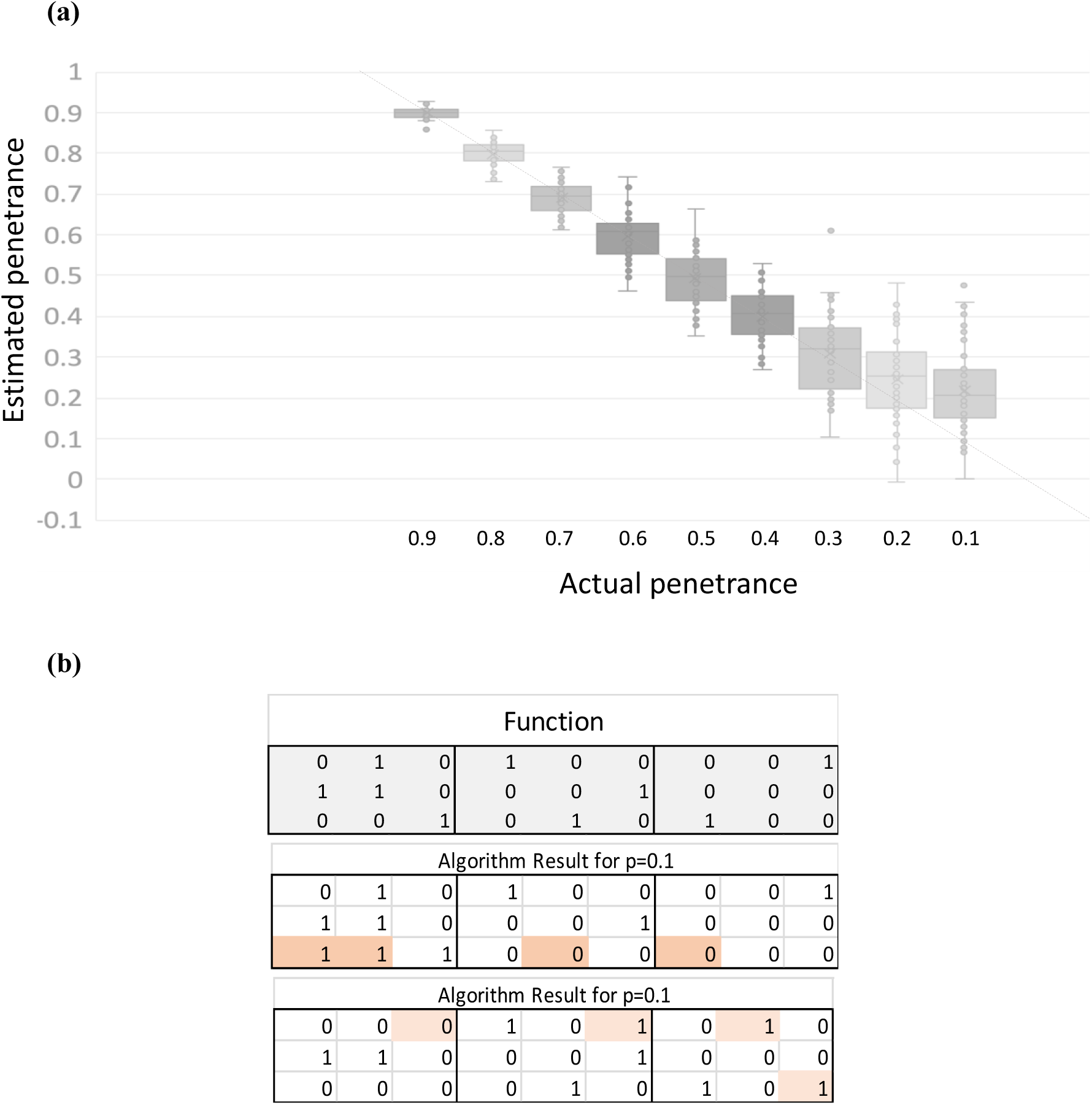
Analysis of simulated data. **(a)** These are values of the penetrance calculated for 50 simulated data sets each for 9 values of penetrance, *p*. For all of these values, except the two rightmost (*p* =0.2 and *p* =0.1), the greedy algorithm returned the exact function. **(b)** For these two there were a few errors in the function (top panel is correct function), as shown in these examples for two cases of *p* =0.1 simulations (the errors are highlighted).

We generated the genotypes randomly, making the simplifying assumption of equal allele frequencies, and used this function to determine the phenotypes, then added uniform noise to the tensor using the relation of Equation 19 to generate the data set for specific values for the penetrance. Correction for allele frequencies is a simple linear transformation. The algorithm was used to infer the discrete function and estimate the penetrance. The results, both for this function and others not shown here, demonstrated that the algorithm works well to infer the exactly correct discrete function for all values of the penetrance greater than about 0.24. Penetrance levels less than this value (high “noise” levels) lead to some incorrect entries, as shown in Figure 5.

It is clear from these results that the simple algorithm provides a reasonably robust method for inferring a complex discrete function from data as well as estimating the penetrance. For larger data sets, of course, the thresholds for inference errors will be smaller than seen here.

### 5.7 Genetic heritability

The quantitation of heritability has been an important, long-standing problem in quantitative genetics originally approached by consideration of variance. The ideas of what are called broad and narrow sense heritability have their roots in the Fisher paradigm [1,2]. In the classical model the components of the trait value (phenotype) are commonly these: the population mean, the genetic effect and the “error term”. The assumption of normally distributed components, with no covariance imposes a model-dependence on the analysis. This model is convenient in that it implies that the variances add, but its validity is often in question. The broad sense heritability is simply defined as the ratio of the genetic variance to the population average, or the fraction of trait variance that is due to all genetic. In this connection it is important to remind ourselves that Huang and Mackay have clearly shown that contributions of the additive, epistatic and dominance variance components in the classical descriptions do not contribute only to these three respective variance components [4]. This means then that the variance components cannot be attributed to these model components, which has often been done in genetic analyses [4]. Genomic heritability is the fraction of the genetic variance that can be explained by regression on the markers and will only be quantitatively accurate when the genotypes of all causal variants are known. See [30] for a useful further discussion of heritability for panmictic populations.

When there is a way to determine the additive component of the genetic variance from separate experimental data, the fraction of the trait variance that is attributed to the additive variance is the classically defined “narrow sense” heritability. Classical methods often use the analysis of variance of full and half-sibling families and use maximum-likelihood methods for relatives with different degrees of relatedness to estimate this quantity.

In the information theory formalism heritability, in the “broad sense,” can be reduced to a quantity that is actually rather simple to state. Since the total dependence, including all components additive and non-additive alike, between a set of loci and a phenotype can be quantitated by the multi-information, we can use this quantity effectively to define heritability if we have calculated the penetrance for each set of loci. It is important to emphasize that the total dependence, measured by the multi-information, includes all effects, both additive and nonadditive effects. For a given phenotype then we propose to define the heritability as the ratio of the total of all the dependence (this means the multi-information for all subsets of dependent variables affecting the phenotype), times the respective penetrance for each subset, divided by the “maximum possible” dependence for the same sets of loci. The maximum possible dependence is clearly the multi-information assuming full penetrance for all dependencies, and therefore no effective “noise”. Therefore, the heritability, 𝓗 can be written by the expression

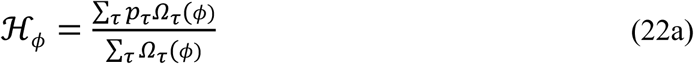

where *τ* is a subset of variables containing the phenotype variable and genetic loci. These are the sums over all subsets of the full set of all genetic loci exhibiting dependence for the specific phenotype, *ϕ*. In the numerator with the corresponding penetrances and in the denominator without. If the penetrance is full for all determinants of the phenotype the heritability reduces to one – full heritability. This means that there are no environmental effects, unaccounted for loci or subsets of variants, or other sources of “noise”. In order for this to be a valid heritability for trait *ϕ*, of course, all possible genetic variable subsets must be included for which *Ω*_*τ*_(*ϕ*) is non-zero. Equation 22a is then a valid and rigorous abstraction, but one that requires all possible multi-locus effects to be quantitated to actually calculate.

One might think that since the sum over dependent tuples may not be disjoint, having some overlaps, that the dependency could be overcounted. In other words, some loci may participate in several subsets of dependent loci. This kind of potential overcounting is not a problem, however, since the measures are weighted by the penetrance and normalized by the total sum, including all possible overlaps. Note that in this definition of heredity we need not assume linkage equilibrium. In fact, Equation 22a applies in the completely general case, with or without linkage disequilibrium.

Practically we must limit the sum to those subsets of variables whose dependencies are detectable and significant, so the criteria for significance must also enter the determination of heritability. This is not because weak dependencies don’t count, but because the calculation of *p*_*τ*_ can only be accurate if the dependence is significant. This definition is different from the classical description then in yet another way. We are calculating the heritability of traits based on all the variants considered in the analysis, while the variance form purports to include all genetic effects but is dependent on the unknown range of genetic differences in the population considered.

If the dependencies for a phenotype were all single locus effects (pairwise dependence) then the heritability would only be a function of the mutual informations between these loci and the phenotype, *ϕ*:

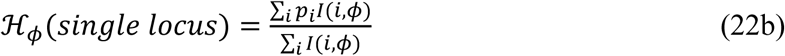

where *I*(*i, ϕ*) is the mutual information between the loci and the phenotype *ϕ*.

In this paper we also restrict the sum to triplets, subsets of three variables, two loci and the phenotype of interest, although Equation 22a is certainly valid for any size of subset *τ*, and any number of genetic loci. Since the composition of dependence for each triplet can be clearly separated into the components due to single locus and two loci dependence, as long as the two loci are independently segregating (*I*(*i, j*) = 0) we can also then separate the heritability into two components by separating the sum in the numerator into two parts, the pairwise or additive effect and the three-way effect. In this case, from Equation 9b it is clear that for the heritability limited to two locus epistatic interactions (no single locus additive effects), 𝓗_*ϕ*_(*triple*), we have

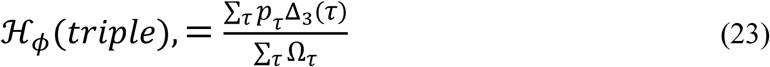

where *τ* indicates all triple dependencies. This formulation provides a rigorous and complete description of heritability given the division between the genetic determinants and the unknowns, the “noise”. It also provides a practical way to calculate the heritability under specific assumptions. Contrast this with Fisher’s heritability, *in the broad sense*, which is the ratio of the variances of phenotype to genotype in the population. Narrow sense heritability is more important in the sense that it quantitates the proportion of the phenotypic variation that is transmitted from parents to offspring [30]. The argument for this interpretation that ignores epistatic effects, which are frequently disrupted by segregation, is plausible, but it is certainly incomplete.

### 5.8 Protective alleles: Interactions that nullify effects

The interaction of two loci, of course, means that each locus may modify the effect of the other in some way. In medical genetics it is becoming increasingly clear that there are potentially severe effects of pathological variants that are not realized in phenotypes. This means that the genetic background of single or multiple loci is providing an effect that “protects” the subject from the pathology. This is an important area of research at the moment. These recent examples for Alzheimer’s disease are emblematic of the approach [31,32] which promises to provide biological insights into the mechanisms of pathology, therefore the nature of the genetic functions expected to be encountered in analysis of genetic data are worth investigating. What is clear from our formulation is that certain discrete functions involving two loci can exhibit a protective-like character which can be characterized. To make this more precise, and to illustrate this kind of interaction in our formalism, we look at a specific, concrete case to examine the instance of protective alleles. The hallmark of protective effects is easily described in terms of the genotype-phenotype table. For simplicity let us consider a binary phenotype where 1 is a negative phenotype, a disease state, and 0 is normal. The effect of a protective allele then simply means that one variant of gene A has the effect of reversing the disease effect of gene B and making the phenotype normal. This can be viewed as a kind of dominance, but a simple model example illustrates the point. A model that shows a protective effect given the gene assignment above is illustrated in Table 3.

**Table 3.**
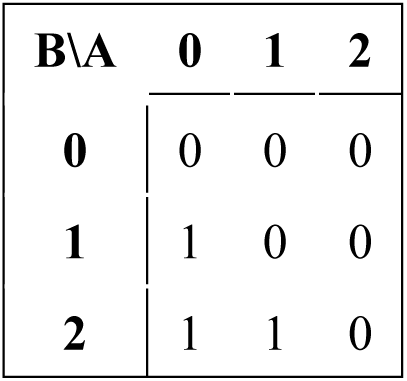
A genotype-phenotype table (100% penetrance) illustrating a protective effect of gene A alleles on the disease-causing alleles of gene B. Phenotype 1 indicates diseased, 0 is normal.

The minor allele of gene B (one or two copies) is assumed to cause the pathology except that it is neutralized in the presence of the minor allele of gene A (one or two copies), which is the protective allele. There are, of course other functions that exhibit such effects.

In order to illustrate the systematic effect in a very simple case we consider the haploid genetics case with binary phenotypes, where there are only 16 possible genetic models (Figure 6). Only four of the 16 possible 2×2 genetic models exhibit protective effects.

**Figure 6.**
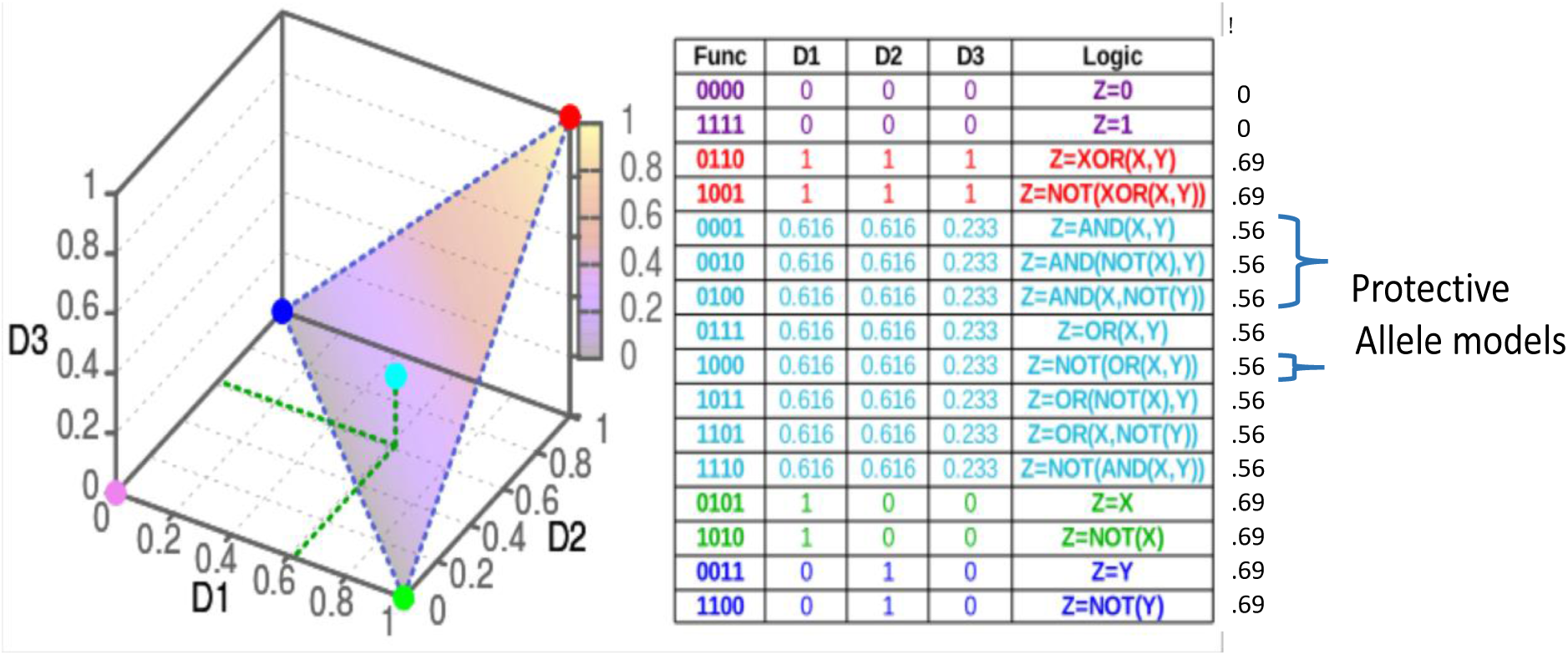
Four of the 16 2×2 genetic models show protective effects. The functions are shown in linear form and color coded according to the families as marked on the information plane.

## 6. Analyzing a Yeast Genetic Data Set

To illustrate the application of the information theory approach to quantitative genetics we analyze a data set of haploid data from a large yeast cross generated by Kruglyak and colleagues [33]. The data consist of 4,390 haploid strains resulting from the cross of a wild, vineyard strain, and a widely used laboratory strain of *Saccharomyces cerevisiae.* This is an F2 cross, so that the recombinations between the two parental chromosomes occur in a single meiosis event for each of the resulting strains. The resulting haploid strains are essentially the gametes from the hybrid F1 strains. The data includes genotypes of all 4,390 strains, at 28,820 SNP positions, and 20 phenotypes, average growth rates under different conditions and in the presence of different compounds. We have restricted our use of the data to those phenotypes that showed a relatively high reproducibility in replicates. We used only those phenotypes whose replicates exhibited highly consistent correlation coefficients. These criteria, a replicate correlation coefficient above 0.8, selected 4 of the 20 phenotypes reported by Bloom, *et al*. [33]. We report the analysis of two of these four phenotypes: growth in the presence of Neomycin (correlation coefficient 0.86), and Copper sulfate (0.82).

### 6.1 Genetic dependencies

We calculated the pairwise effects, mutual information, between single genetic variants and the phenotype, and the measure of three-way effects, using a representative set of 100 variant markers across the genome. To calculate the three-way interactions accurately we wanted independently segregating markers, so we selected a set of 100 markers that were isolated by iteratively eliminating one of each pair of markers that had a mutual information of more than 0.05. The markers were widely spread, and we calculated the recombination frequencies between each pair of neighboring markers to assess the statistics of segregation. The results for pairwise and three-way genetic dependencies for the two phenotypes are shown in Figures 7-10. For our significance calculations, we follow the permutation strategy proposed by Churchill and Doerge [34]: we shuffle the input data, breaking the connections between genetic markers and phenotypes, compute the dependency scores of all shuffled tuples, and count how many randomized scores are above the original score of interest. We repeat this procedure 100 times tallying the number of scores above the score of interest. The *p*-value is then the fraction the exceeding randomized scores take in the total number of tuples times 100.

**Figure 7.**
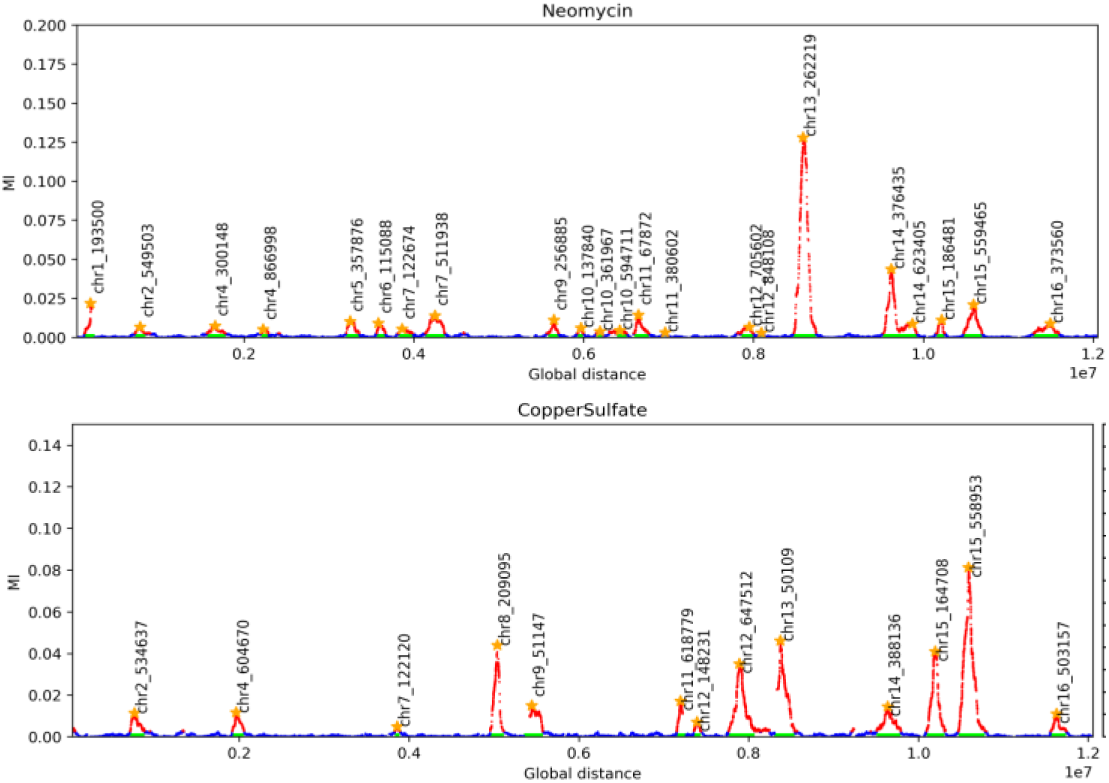
The pairwise peaks for the genetic determinants of two phenotypes. The locations indicated are the chromosomal coordinates of the highest scoring marker in the peak.

The two panels in Figure 7 show the pairwise peaks resulting from plotting the mutual information for the entire set of 28,820 genetic markers for each of the two phenotypes. The entries in Table 4 show the location of the highpoint of each peak, the standard deviation and the width. The width of the peak is defined by the outermost boundary of the peak determined by the locations of the last significant marker by mutual information on each side of the peak. The peak widths in this case are largely due to the co-segregation of contiguous blocks in the F2 meiosis.

**Table 4.**
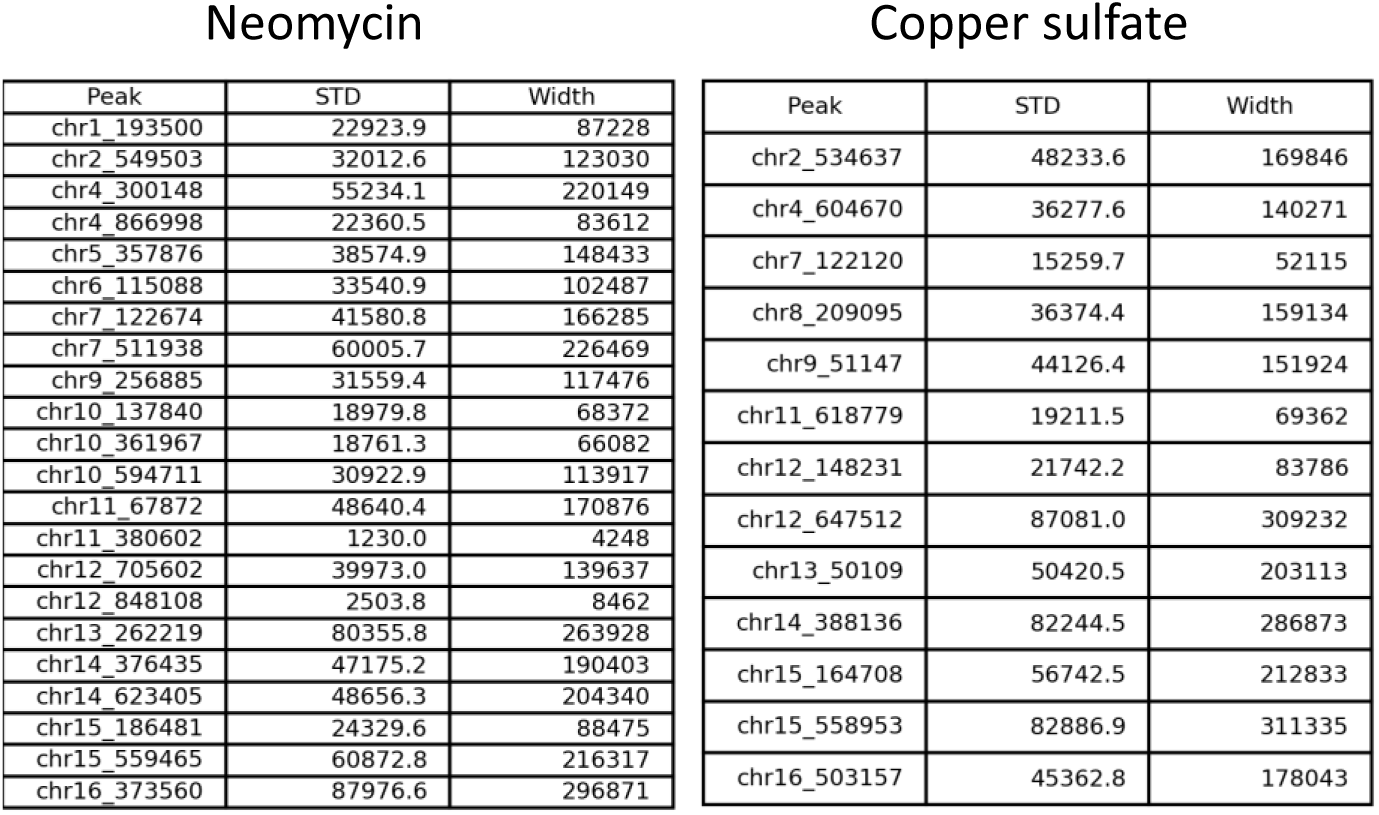
Pairwise peaks: global location, standard deviation and widths. The coordinates here are the global coordinates in base pairs, beginning with chromosome I. The widths and the standard deviations are in base pairs. The width is the distance between the two extreme, significantly scoring markers.

| Neomycin |  |  | Copper sulfate |  |  |
| --- | --- | --- | --- | --- | --- |
| Peak | STD | Width | Peak | STD | Width |
| chr1_193500 | 22923.9 | 87228 | chr2_534637 | 48233.6 | 169846 |
| chr2_549503 | 32012.6 | 123030 | chr4_604670 | 36277.6 | 140271 |
| chr4_300148 | 55234.1 | 220149 | chr7_122120 | 15259.7 | 52115 |
| chr4_866998 | 22360.5 | 83612 | chr8_209095 | 36374.4 | 159134 |
| chr5_357876 | 38574.9 | 148433 | chr9_51147 | 44126.4 | 151924 |
| chr6_115088 | 33540.9 | 102487 | chr11_618779 | 19211.5 | 69362 |
| chr7_122674 | 41580.8 | 166285 | chr12_148231 | 21742.2 | 83786 |
| chr7_511938 | 60005.7 | 226469 | chr12_647512 | 87081.0 | 309232 |
| chr9_256885 | 31559.4 | 117476 | chr13_50109 | 50420.5 | 203113 |
| chr10_137840 | 18979.8 | 68372 | chr14_388136 | 82244.5 | 286873 |
| chr10_361967 | 18761.3 | 66082 | chr15_164708 | 56742.5 | 212833 |
| chr10_594711 | 30922.9 | 113917 | chr15_558953 | 82886.9 | 311335 |
| chr11_67872 | 48640.4 | 170876 | chr16_503157 | 45362.8 | 178043 |
| chr11_380602 | 1230.0 | 4248 |  |  |  |
| chr12_705602 | 39973.0 | 139637 |  |  |  |
| chr12_848108 | 2503.8 | 8462 |  |  |  |
| chr13_262219 | 80355.8 | 263928 |  |  |  |
| chr14_376435 | 47175.2 | 190403 |  |  |  |
| chr14_623405 | 48656.3 | 204340 |  |  |  |
| chr15_186481 | 24329.6 | 88475 |  |  |  |
| chr15_559465 | 60872.8 | 216317 |  |  |  |
| chr16_373560 | 87976.6 | 296871 |  |  |  |

In Appendix D, supplementary Tables D1 and D2 we indicate the quantitative results for the inter-chromosomal interactions shown in Figure 8. We decided to omit intra-chromosomal interactions because in this cross with only one meiotic recombination the correlated blocks of markers are significant and even the widely spaced markers used here have some residual correlation. The inter-chromosomal pairs have no such systematic correlations because of the nature of the cross, and thus the calculations of their epistatic effects are most accurate, as explained above. The fractions of the interactions that are attributed to epistatic and additive effects are indicated in the final two columns of the supplementary tables. There are variable levels of epistasis evident, but they are all below 10%.

**Figure 8.**
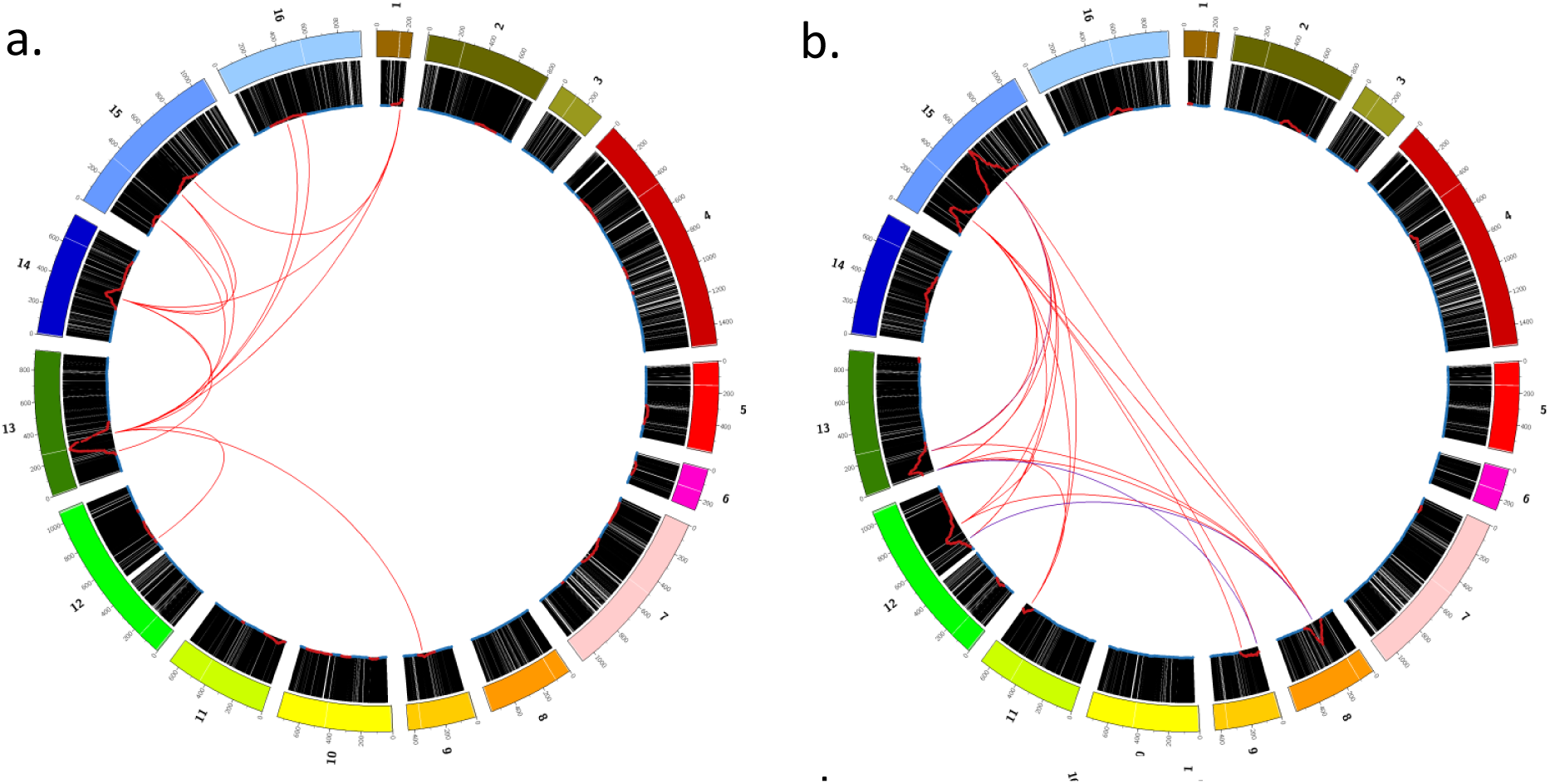
Genetic effects for two growth phenotypes. The full genome is shown with the single and two locus effects. The pairwise peaks for these phenotypes, as shown in Figure 7, are indicated as the red curves in the black band (using all 28,820 markers.) The variant pair interaction effects for these phenotypes are indicated by the internal red lines (all interacting pairs are shown in the supplementary tables, D1 and D2) indicating the significant three-way dependencies between the two markers at the ends of the line and the phenotype, indicating genetic loci interacting. **a.** Genetics of Growth on Neomycin, **b.** Growth on Copper sulfate.

A gene known to affect copper resistance, the well-known metallothionein gene, CUP1, which is in the peak on chromosome 8, participates in several interactions. Note for all the interactions described here that since the markers used for this analysis have relatively low resolution, two neighboring markers may well indicate the same epistatic interaction. This is particularly likely when these markers are within the same mutual information peak, but we do not attempt to separate these effects here.

There are several notable features shown in the data of Figure 9, which compares the epistatic fractions for the two phenotypes. These quantities are the fractions of the total dependence of each of the two genetic loci that are attributed to their interaction. Note primarily that the epistatic fractions vary several-fold, and the variation among them includes interacting pairs with the same marker, so that the degree of epistasis is clearly determined by both markers. The left-most six interactions of the copper sulfate panel, for example, indicates the interactions with the marker on chromosome VIII near the CUP1 locus.

**Figure 9.**
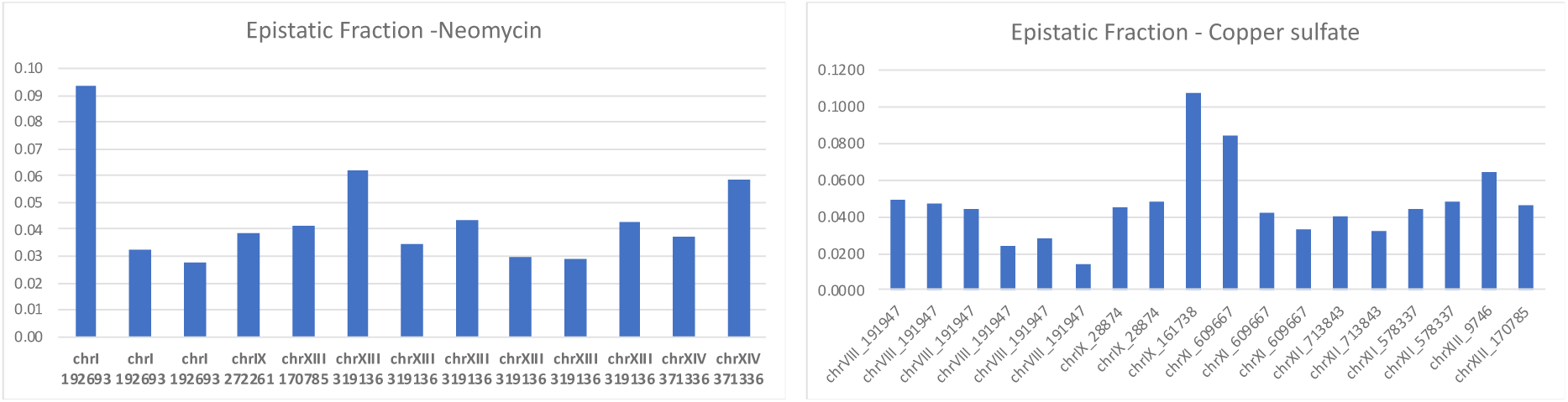
Epistatic fractions compared. The bars represent the fractional epistatic effect for the interactions in the order listed in supplementary Tables S5a and S5b. For brevity the marker indicated below each bar is the one listed in the left-hand column of the tables, and represents the pair.

To see an example of the detected epistatic interactions in the data, we analyzed the tuple which had the largest detected multi-information, Ω, with the Neomycin phenotype: loci chrI_319136 and chrXIV_371336. As shown in Table D1 this tuple has a multi-information of Ω = 0.1116, and an epistatic fraction of only 0.062 (*i.e.*, an additive fraction of 0.938). The phenotype distribution for samples of each unique genotype is shown in Figure 10. Even with this large additive fraction, it is clear from the phenotype distributions that we cannot consider this interaction to be entirely additive. Genotype 10 has the highest median phenotype at 0.624. Flipping either locus to 00 or 11 results in medians of 0.157 and 0.234, respectively (decreases of 0.467 and 0.39). If these effects simply added, we would expect a decrease of about 0.86. The median effect on phenotype 01, however, is a decrease of nearly 1.4, well beyond what would be expected from an additive effect. This is corroborated by a qualitative assessment of each distribution: the distribution of genotype 01 is markedly different from that of any other genotype, implying that even the relatively small epistatic interactions which we detect have a real and noticeable consequence.

**Figure 10.**
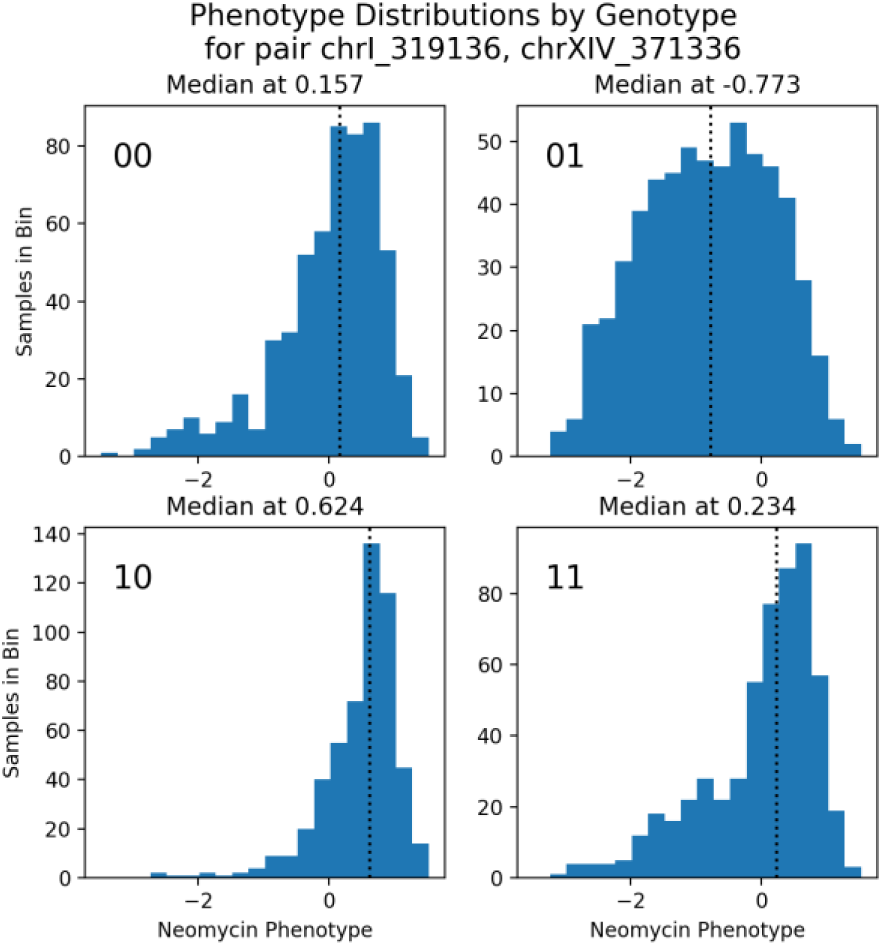
Phenotype distributions by genotype. We examined the tuple with the highest value of Ω for the Neomycin phenotype: loci chrI_319136 and chrXIV_371336. The panels show the phenotype distribution for each genotype (*e.g.*, plot 01 shows samples with chrI_319136=0 and chrXIV_371336=1).

### 6.2 Pleiotropic effects

Recall that pleiotropy is the effect wherein a single genetic locus causes multiple phenotypic effects. For three-way dependencies this means a triple set of variables showing dependency that consists of a single genetic locus and two phenotypic variables. In the yeast data set we are considering this is reflected in a pattern that is easy to observe. Consider Figure 8 and ask what a pleiotropic effect would imply. One instance would be, for example, a situation in which a single pairwise dependence is evident in both the Neomycin and the Copper Sulfate resistant phenotypes. There are actually several such cases, but the most evident one that will use as an example are the two single locus peaks that appear on chromosome XV in the figures showing both phenotypes, Figures 8 a and b. The tables in Appendix D illustrate the pleiotropy more specifically, identifying these two different variants that are co-identified for both phenotypes: these markers: chrXV 605280, and chrXV 189231. If we carry out the corresponding triple dependence measure using the symmetric delta, we do see these same loci showing up (results not shown). The symmetric delta is, of course, applicable for any combination of variables to detect collective dependence and the occurrence of pleiotropy, among two phenotypes and one genetic variant, is a collective dependence just as is the dependence among two genetic loci and one phenotype.

## 7. Discussion

With the formulation presented here we have begun to build a structure based on information theory for describing and analyzing data in quantitative genetics. The advantages of an information theory approach to quantitative genetics are multiple and include these: a model-free, agnostic approach to multivariable dependence, the simplicity of the formalism, and the fundamental separation of dependence detection from the ascertainment of the functional nature of the dependency. There are important differences from the classical formulation: (1) GWAS includes a tacit assumption of linearity (the usual association studies largely make use of correlation coefficients, which by their nature quantify linear dependence, and thereby tacitly assume a linear model), and (2) variance component analysis assumes other tacit model assumptions which are discussed in the introduction. These differences from the classical formulation permit us to formulate important genetic quantities in a direct, simple and calculable way. This contrasts with the classical formulation, for example with respect to the variances of phenotypes and genetic variance and the additivity of variances on the assumption of Gaussian distributions, and other more subtle assumptions point out by Huang and Mackay [4]. The assumption about which components contribute exclusively to which variances (additive, dominant, and epistatic) are based on models which are generally not valid, and this flaw renders the inference of genetic architecture from variance component analysis invalid [4]. The conclusion is summarized in this paper as follows: “The crux of the problem is the undesirable features of the classical model as well as the alternative parameterizations that there is not a one-to-one correspondence between gene action at underlying quantitative trait loci and the partitioning of variance components except under very specific and restrictive circumstances”. Many researchers have pointed to the need for an alternative approach to the classical model [3,4] and our paper begins the construction of such a formulation using information theory. We fully recognize that the formulation in this paper, limited to panmictic populations, provides only the first steps and that more development, which we plan for future publications and encourage others to address, is definitely needed. It has been well remarked that no natural population is panmictic, and that linkage disequilibrium, relatedness and population structure must also be described in the information theory formulation. These issues are amenable to an information formulation, while some of them can be somewhat more complex. They have been approached in the use of information theory for discussions of evolution and populations previously [28,29], and we will treat these additional issues in a future publication.

Since the formulation presented here uses discrete functions extensively it is important to understand the advantages and limitations of this underlying structure. While there are a finite number of discrete functions for any finite number of variables and alphabets, there are an infinite number of possible (discrete plus continuous) distributions for any finite number of variables and alphabets. For example, there are 3^9^ =19,683 3×3 discrete functions, (3 variables and three-letter alphabets). This obvious distinction is important, and it is clear that the function-to-distribution mapping is not one-to-one – there are vastly more distributions than discrete functions. Since we have assumed no linkage disequilibrium for the most part in this paper, the “information landscape” for two variants was confined to a plane. When the genetic variants are not independent variables, and linkage is present, the “information landscape” is no longer a plane and requires a more complex description.

It is clear that the addition of “noise” to the discrete functions (see [19] and section 5.1 generates distributions around the discrete functions at varying distances in the information landscape. The point where the uniform noise distribution fully dominates and masks all information content, the “black hole” of information [19], represents the single distribution with uniform probabilities – it has maximal entropies. It may seem extraordinary, however, that this point has finite and distinct coordinates in the information landscape, as we describe in section 5.2. This points to the fact that the landscape includes functions with a wide range of penetrance and heritability, and that they are not at all distributed continuously on the landscape.

What is important here is to clearly define the distinctions and classifications that information theory can provide, and what these measures do and do not distinguish. This is an important specific question. One striking example is that the distribution of the “black hole” [19] and the discrete function that is XOR-like (with no non-zero pairwise measures) have distinct and unique coordinates in the information space for three variables, but as the alphabet size increases without bound (allowing more and more precise definitions of the variable values) they converge in the normalized space coordinates. It is a notable and related fact that the maximization of the symmetric delta leads to exclusively three-way dependent functional relationships, as shown in Appendix B.

The application of this information-based formalism to genetics brings forward a number of interesting and important relations. The quantitation of the penetrance, for example, is simple and direct, but depends on a very clear division between what variables are being included to make the inferences and what is being ignored and therefore being considered to contribute to the “noise”. We characterize the non-included variables with the “noise,” even if they include the more complex genetic interaction effects which are not included. This is obvious and commonplace, but our formulation forces it to be explicit in all cases. Likewise, the information theory expression for heritability is both direct and intuitive, but also depends, as it must, on the precise assumptions made, and on the significance of the dependencies. Here we have ignored any variant interactions among more than two genetic loci, and effects that involve three or more loci can therefore contribute noise directly to the penetrance and heritability calculations. We are thus forced to be explicit about what is meant by genetic background effects, which also include any effects of variants not considered as variables in the analysis.

Since the yeast two-strain cross involves only a single meiosis per strain to produce the collection analyzed, individual marker segregation within chromosomes is not a good assumption since blocks of markers will necessarily segregate together, while individual chromosome segregation, on the other hand, is generally an excellent assumption, and is supported by the data. In future work we will consider the more complex cases where the possibility of non-independent segregation, and non-uniform population structures, are included, as in human data. The extension of the formalism to quantitate these effects and their implications will be an important part of the full description.

Bloom, *et al*. [33] argue that the additive effects are much greater than the epistatic effects on the quantitative traits they measured using the variance component method, estimating a 9% overall interaction effect. Our results agree with this qualitative conclusion, but we calculate the interactions specifically for each pair of inter-chromosomal markers and found that the levels of significant interactions among variants are highly variable, but in this general range. Huang and MacKay specifically point out in [4] that Bloom, *et al.* used the variance component calculations improperly to make this estimate. It is therefore not surprising to find quantitative disagreement with our results, even if the general quantitative ranges are similar. Results in similar F2 crosses in mice also show small epistatic effects as we would expect [40]. These authors use logistic regression methods to look at both pleiotropic and epistatic effects of pairwise identified loci.

We argue that the ability to quantitate the pairwise, additive and epistatic effects unambiguously is definitely a significant step forward and provides a distinctly different and practical alternative to the classical model. Comparison of our specific results with those of Bloom, *et al*. show close agreement for the pairwise effects, but unfortunately there is no way to make the full comparison for multi-loci effects for the overall estimate of additivity. The major difference with [33] is that we accurately calculate the specific fraction of additive and epistatic effects for each pair of loci.

The present contribution, a first step towards a full information theory of quantitative genetics, leaves a number of important problems unaddressed. These include the characterization of population structure, relatedness and linkage disequilibrium, as we have mentioned. Their incorporation into the formalism, so they are accounted for in the genetic inferences, will be an important improvement. Genes with significant linkage may also interact. This is often a problem in genetic analysis, the disentangling of interaction from disequilibrium, and the information theory formalism method needs to be extended to treat these cases accurately. While larger data set sizes are required to accurately assess more interacting loci than two the formalism can certainly address these more complex dependencies, which will be more and more important in future as data sets of genetic information continue to grow rapidly. Future work will focus on addressing all these issues and incorporating extensions into a formalism that will provide a broadly applicable set of descriptive and analytic tools.

## Acknowledgements

This work was made possible by partially support from the NSF National Science Foundation, EAGER grant (IIS-1340619), by the Pacific Northwest Research Institute, and the Bill and Melinda Gates Foundation. In the early stages of this work the Luxembourg Centre for Systems Biomedicine (LCSB) of the University of Luxembourg also provided significant partial support. We thank Joe Nadeau, Aimee Dudley for stimulating discussions, Greg Carter and Philip Welkoff for useful comments on an early version of the manuscript, and Elaine Skeffington for expert copy and format editing of the manuscript. None of the views expressed here should be attributed to the funding organizations. Neither did they play any role in the decision to publish or in the process of publication.

## Author contributions

DG conceived of the project, DG, NS and JK-G provided and developed key ideas for specific parts of the formalism and paper sections. LU and JK-G carried out calculations on the yeast data. DG wrote the first draft of the paper and DG, NS and JK-G further edited and revised the manuscript.

## Supplementary Materials - Appendices

### Appendix A. Two propositions regarding key inequalities

#### Proposition 1

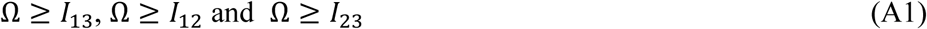

Proof:

We first prove that Ω ≥ *I*_13_.

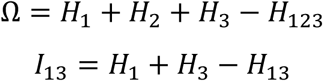

Subtract the lower from the upper, to get Ω − *I*_13_ = *H*_2_ − *H*_123_ + *H*_13_. Since Ω ≥ 0, and H_*i*_ ≥ 0, and since the sum of the entropies of any subset of variables is greater than or equal to the joint entropy, it is clear that *H*_2_ + *H*_13_ ≥ *H*_123_, and the proposition is proven. Starting with *I*_23_ or *I*_12_ yields the parallel inequalities. There are, however, tighter bounds.

#### Proposition 2

If Ω_123_ is the multi-information for three variables, then

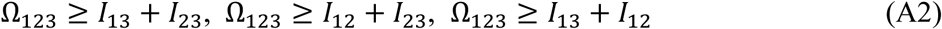

By definition Ω = *H*_1_ + *H*_2_ + *H*_3_ − *H*_123_ and the mutual informations, and interaction information, *I*_123_, are

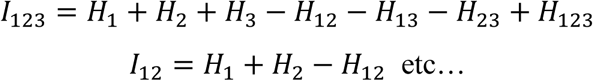

So we have

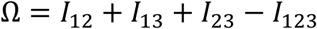

Since the recursion relations for the interaction information are

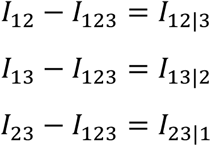

we can write

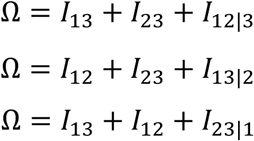

Since by the definition of the mutual information *I*_12|3_ = *H*_1|3_ + *H*_2|3_ − *H*_12|3_, and we have the identity *H*_2|3_ + *H*_12|3_ = *H*_1|23_ by the basic probability identities. Since *I*_12|3_ = *H*_1|3_ − *H*_1|23_ and

*H*_1|3_ ≥ *H*_1|23_, *I*_12|3_ ≥ 0, the above three equations imply,

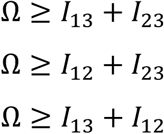

And the proposition is proved.

### Appendix B. Maximizing the Symmetric Delta finds any XOR-like dependence first

Equation 9a implies that if the mutual informations are all zero or very small, for a triplet, the deltas must all be equal, or close to it. Furthermore, they must all be equal (or close to it for small mutual informations) to the multi-information, Ω. The product of the deltas, the symmetric delta, is maximized when the factors are equal given a constant sum. This means that maximizing the symmetric delta will tend toward picking out the functions with deltas that are closest, given that the total correlation or multi-informations, Ω, are constant. The XOR-like functions, which have no pairwise, but do have three-way dependence, therefore maximize the symmetric delta for given Ω. An easy calculation shows the maximum of the symmetric delta, 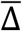, occurs when all factors are the same. If we define variable measures of the pair-wise dependence, {*x*_*i*_} then

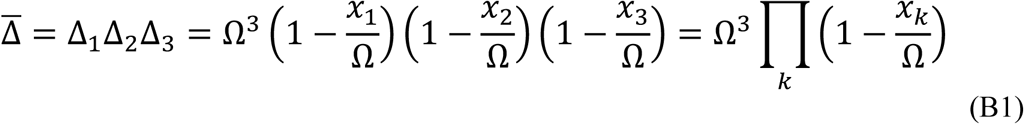

where

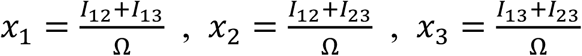

For all triplets, indicated by *k* and *l*, we then can say that 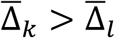 only if

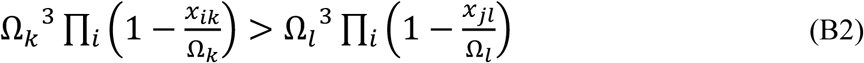

It is clear from this equation that for similar values of Ω the largest is the one with all x’s equal to zero, which is the XOR-like case. Thus, if there is an XOR-like dependence present in the data it will be at the top of the symmetric delta scoring list.

### Appendix C. A matrix formulation of the 3-way dependence relations

The set of equations in 9a can be expressed simply in matrix form by defining the vectors:

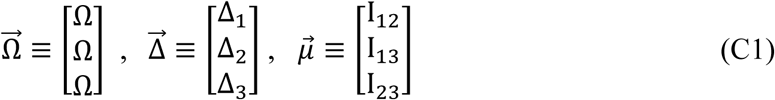

All of the components of these vectors are non-negative, and for the moment we are not assuming that *I*_12_ = 0, so this holds for the general case. Then we have a matrix relation,

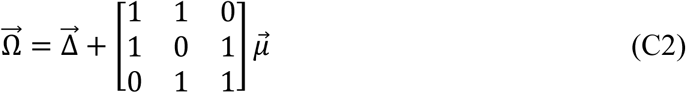

The functions are thus confined to the landscape in only the non-negative sector (all coordinates are ≥ 0.

The matrix 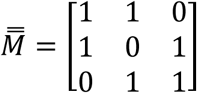, has the inverse, 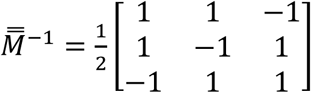, and writing the expressions for the other two vectors gives these simple equations

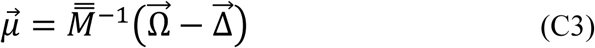

Thus, the 3-variable relations reduce to a simple equation in matrix form.

### Appendix D. Interactions in the yeast data [33]

**Table D1.** **The inter-chromosomal interactions for the Neomycin phenotype**, exhibiting the markers (identical markers are color-coded for better visualization. The p-values, calculated from the permutation of the data, are approximately the same.

|  | Marker1 | Marker2 | Omega | p-value | Epistatic Fraction | Additive fraction |
| --- | --- | --- | --- | --- | --- | --- |
| I-XV | chrI 192693 | chrXV_605280 | 0.0345 | 2.020E-06 | 0.0936 | 0.9064 |
| I-XIII | chrI 192693 | chrXIII_319136 | 0.0856 | 2.020E-06 | 0.0324 | 0.9676 |
| I-XIV | chrI 192693 | chrXIV_371336 | 0.0633 | 2.020E-06 | 0.0275 | 0.9725 |
| IX-XIII | chrIX 272261 | chrXIII_319136 | 0.0718 | 2.020E-06 | 0.0385 | 0.9615 |
| XIII-XIV | chrXIII 170785 | chrXIV_371336 | 0.0594 | 2.020E-06 | 0.0415 | 0.9585 |
| XIII-XIV | chrXIII 319136 | chrXIV_371336 | 0.1116 | 2.020E-06 | 0.0620 | 0.9380 |
| XIII-XIV | chrXIII 319136 | chrXV_189231 | 0.0757 | 2.020E-06 | 0.0346 | 0.9654 |
| XIII-XIV | chrXIII 319136 | chrXVI_433340 | 0.0693 | 2.020E-06 | 0.0435 | 0.9565 |
| XIII-XV | chrXIII 319136 | chrXV_455183 | 0.0696 | 2.020E-06 | 0.0297 | 0.9703 |
| XIII-XIV | chrXIII 319136 | chrXVI_304984 | 0.0711 | 2.020E-06 | 0.0290 | 0.9710 |
| XIII-XII | chrXIII 319136 | chrXII_578337 | 0.0686 | 2.020E-06 | 0.0429 | 0.9571 |
| XIV-XV | chrXIV 371336 | chrXV_189231 | 0.0539 | 2.020E-06 | 0.0373 | 0.9626 |
| XIV-XV | chrXIV 371336 | chrXV_455183 | 0.0491 | 2.020E-06 | 0.0582 | 0.9418 |

**Table D2.**
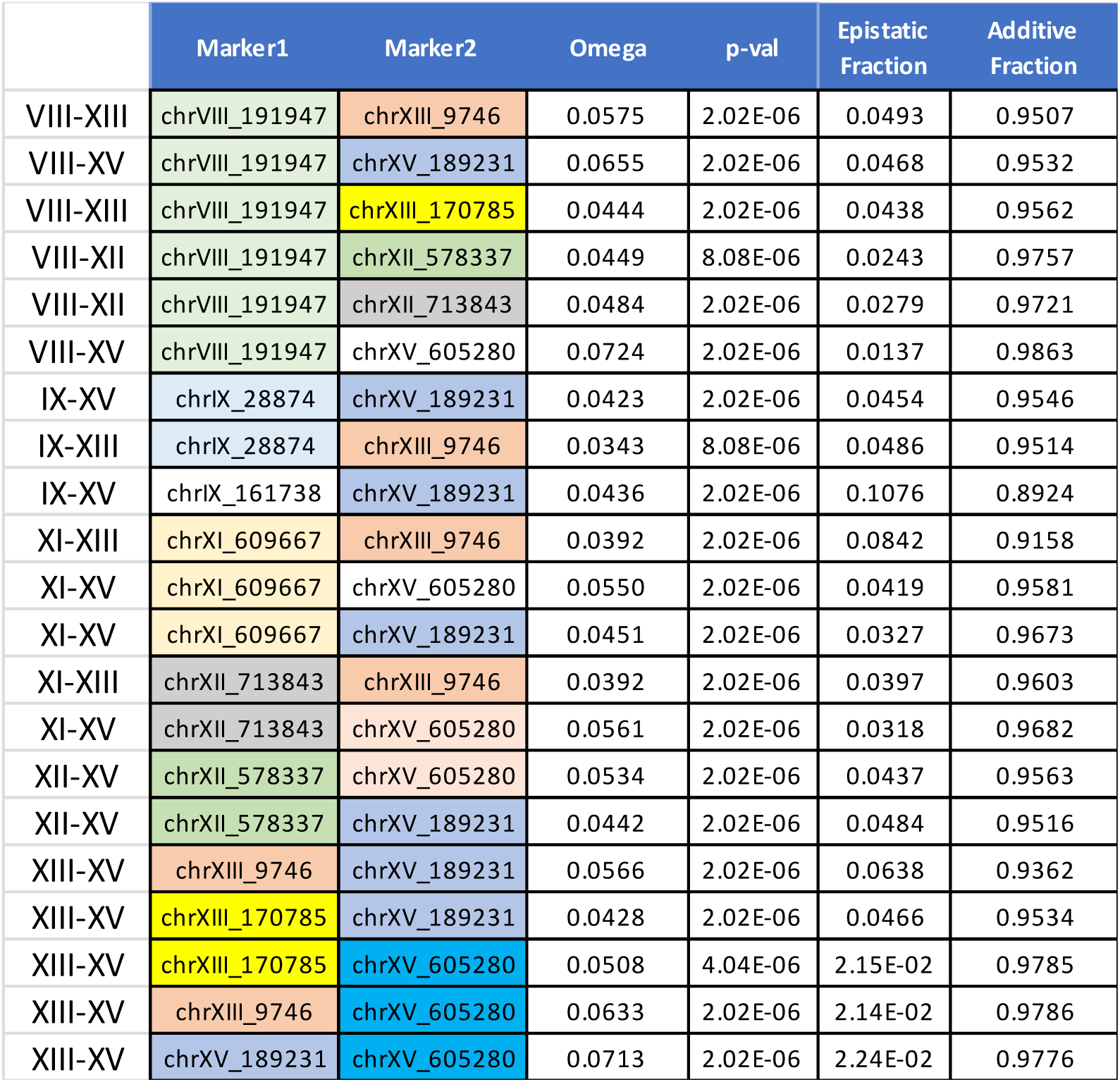
**The inter-chromosomal interactions for the Copper sulfate phenotype**, exhibiting the markers (identical markers are color-coded for better visualization). The p-values, calculated from the permutation of the data, are approximately the same.

### Appendix E. Estimates of entropies from data

There is a large literature on the estimation of entropies and mutual information that dates from shortly after Shannon’s first papers on information theory. This issue needs to be addressed for any application of information theory, including this one, since the data we deal with are used to calculate measures entirely dependent on estimated entropies. It is easy to see intuitively how small sample numbers can bias entropy estimations. Consider that we have discrete categories or alphabets for a variable. If there are only a few samples the principal errors will likely be in underestimating the number of occurrences of categories. This, of course, will cause an underestimate the entropy of the variable. As the sample numbers grow this bias is reduced and the studies have addressed the questions: by how much, possible corrections, the estimation of error distributions etc. [41] A good recent summary of the literature and main issues can be found in [42].

A good feel for the quantitative nature of the estimation problem can be obtained by considering an early formulation of a correction for the estimate of entropy from discrete data. The Miller-Madow bias correction for the maximum likelihood estimator of the entropy of a variable, well discussed in [42], is given by the formula

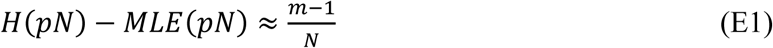

where *H*(*pN*) is the true entropy, *MLE*(*pN*) is the maximum likelihood estimate, *m* the number of letters in the alphabet with nonzero occurrence in the data, and *N* is the total number of samples for the variable. While this correction is not exact, and there is much more to the general problem, it points to the central issue which is the degree to which the data properly represents the distribution of the variable. In our case where the alphabet size is small (for a diploid genotype genetic variable *m*=3) and *N* is orders of magnitude greater, the Miller-Madow correction is therefore very small. This means, in general that the entropy estimates are a good reflection of the true entropies using, as we do here, a simple, so-called naïve, estimate of the probabilities to calculate the entropies. Nonetheless, if the sample numbers decrease or the phenotype resolution increases in some cases, we need to be very aware of the errors in the estimation of the entropies as they affect all information measures.

## Footnotes

1 If *I*_12_ ≠ 0 the situation is more complex, and distinctions need to be made between the redundant information provided by X and Y, the unique information provided, the synergistic information and the quantities indicated in Figure 2 do not apply. This is essentially the information decomposition problem, which has no universally accepted method of computation [39]. We deal with this in a future publication.

